# Genomic analysis of patient-derived xenograft models reveals intra-tumor heterogeneity in endometrial cancer and can predict tumor growth inhibition with talazoparib

**DOI:** 10.1101/2021.03.30.436914

**Authors:** Vanessa F. Bonazzi, Olga Kondrashova, Deborah Smith, Katia Nones, Asmerom T. Sengal, Robert Ju, Leisl M. Packer, Lambros T. Koufariotis, Stephen H. Kazakoff, Aimee L. Davidson, Priya Ramarao-Milne, Vanessa Lakis, Felicity Newell, Rebecca Rogers, Claire Davies, James Nicklin, Andrea Garrett, Naven Chetty, Lewis Perrin, John V. Pearson, Ann-Marie Patch, Nicola Waddell, Pamela Pollock

## Abstract

**Background:** Endometrial cancer (EC) is a major gynecological cancer with increasing incidence. It comprised of four molecular subtypes with differing etiology, prognoses, and response to chemotherapy. In the future, clinical trials testing new single agents or combination therapies will be targeted to the molecular subtype most likely to respond. Pre-clinical models that faithfully represent the molecular subtypes of EC are urgently needed, we sought to develop and characterize a panel of novel EC patient-derived xenograft (PDX) models.

**Methods:** Here, we report whole exome or whole genome sequencing of 11 PDX models and the matched primary tumor. Analysis of multiple PDX lineages and passages was performed to study tumor heterogeneity across lineages and/or passages. Based on recent reports of frequent defects in the homologous recombination (HR) pathway in EC, we assessed mutational signatures and HR deficiency scores and correlated these with *in vivo* responses to the PARP inhibitor (PARPi) talazoparib in six PDXs representing the different molecular subtypes of EC.

**Results:** PDX models were successfully generated from all four molecular subtypes of EC and uterine carcinosarcomas, and they recapitulated morphology and the molecular landscape of primary tumors without major genomic drift. We also observed a wide range of inter-tumor and intra-tumor heterogeneity, well captured by different PDX lineages, which could lead to different treatment responses. An *in vivo* response to talazoparib was detected in two p53mut models consistent with stable disease, however both lacked the HR deficiency genomic signature.

**Conclusions:** EC PDX models represent the four molecular subtypes of disease and can capture intra-tumoral heterogeneity of the original primary tumor. PDXs of the p53mut molecular subtype showed sensitivity to PARPi, however, deeper and more durable responses will likely require combination of PARPi with other agents.

## Introduction

Endometrial cancer (EC) is the most common gynecological malignancy in developed countries with increasing annual rates (1). Whilst most ECs are detected early and have good prognosis, patients with metastatic disease (15%) or who relapse after surgery (∼15%) have a median survival of less than 12 months (2).

EC is comprised of multiple histological subtypes, including low and high-grade endometroid, those with serous and clear cell histology and uterine carcinosarcomas. The Cancer Genome Atlas (TCGA) identified four molecular subtypes: *POLE* mutant (excellent prognosis), mismatch repair deficient (MMRd; intermediate prognosis), *TP53* wildtype (p53wt; intermediate prognosis), and *TP53* mutant (p53mut; worst prognosis) (3). Multiple laboratories have confirmed the different prognoses associated with these subtypes, using a combination of surrogate immunohistochemistry stains or loss of heterozygosity (LOH) analyses and limited sequencing (2, 4-7). Genomic studies of uterine carcinosarcomas (UCS), have also revealed the presence of similar subtypes, however, the majority of tumors (∼90%) contain *TP53* mutations and a low tumor mutation burden (TMB) (8-10).

Apart from the recent approval of immune checkpoint inhibitors (ICIs) for MMRd cancers, there has been little development in terms of precision medicine for EC. Surgery, radiotherapy and chemotherapy still remain the main treatment options. In recent years, there has been a growing interest in applying PARP inhibitors (PARPi) for treatment of EC. PARPi have proven to be incredibly effective in cancers with HR deficiency, such as ovarian and breast cancers with *BRCA1/2* mutations. In EC, PARPi sensitivity was originally reported in cell lines with PTEN protein loss identified as a predictive marker (11), however this was later refuted using a larger panel of cell lines (12). PARPi sensitivity in EC has also been associated with loss of MRE11 protein in EC cell lines (13), and mutations in *ARID1A* (which commonly occur in EC) in cell lines from other cancers (14). Recently, a pan-cancer analysis of bi-allelic alterations in HR DNA repair genes revealed ∼15% of ECs have a combination of germline and/or somatic bi-allelic mutations and/or LOH (15). Another study of TCGA data reported that ∼50% of non-endometrioid ECs (∼90% p53mut) show a mutational or copy number signature associated with defective HR (16).

New targeted therapies, such as PARPi, need to be tested in pre-clinical models that accurately recapitulate the molecular characteristics of patient tumors. In EC, there are a few cell lines derived from UCS and serous ECs. Although multiple cell lines from endometrioid ECs carry *TP53* mutations, almost all of these cell lines are also MMRd and show a high TMB suggesting the *TP53* mutations are acquired through culturing, hence these cell lines are not good models for poor prognosis p53mut EC as they do not recapitulate the biology of this subtype. Identification of effective therapies and predictive biomarkers for p53mut ECs requires well-characterized pre-clinical models that recapitulate this molecular subtype. Patient-derived xenografts (PDX) have been previously demonstrated as reliable pre-clinical models for assessing treatment responses, if carefully characterized.

In this study we performed in-depth genomic characterization of EC PDX models to define their suitability as pre-clinical models and predict HR deficiency status by assessing genomic scars. Here, we report *in vivo* responses to the potent PARPi talazoparib in a panel of MMRd and p53 mutant PDX models, and correlate these responses with genomic features.

## Materials and Methods

### Patient samples

All samples were obtained from patients with informed consent and the study has human ethics approval from the Mater Health Services Human Research Ethics Committee (HREC/15/MHS/127), UnitingCare Health Human Research Ethics Committee (1116), Queensland University of Technology (1500000169, 1500000323) and QIMR Berghofer (P3478, P2095). Clinical data including tumor stage, grade, chemotherapy treatment and survival status was collected (Additional File 1:Table S1). Fresh tissue was obtained from patients undergoing surgery for EC and transported to the laboratory on ice in RPMI, 10% FBS. The remainder was fixed in formalin and embedded in paraffin (FFPE). Where possible, a blood sample was also obtained for sequencing analysis.

### Mouse PDX models

PDX establishment and passaging was performed according to animal ethics approvals at TRI (TRI/021/19) and QUT (1900000701). Fresh primary tumors (n=33) were transplanted into immunocompromised Nod Scid Gamma (NSG) mice within 4 hours of surgery. When transplantation could not be performed immediately, tumors were either stored at 4°C overnight in transport media (n=9) or viably frozen (n=11). Each sample was cut into approximately 1-2 mm^3^ pieces and placed on ice in a 1:1 solution of RPMI:Matrigel. Mice were anaesthetized and a single tumor piece was inserted subcutaneously in the subscapular region (2-4 mice). PDX engraftment was then assessed weekly using micro-calipers. Once a tumor reached a volume of ∼750-1000mm^3^, mice were euthanized using CO2 and several 1-2mm^3^ fragments were transplanted subcutaneously into the next generation of mice. A slice of each PDX was preserved as FFPE, as well as frozen for DNA extraction. Haematoxylin and eosin (H&E) slides were examined by an anatomical pathologist to determine the histology of each PDX passage and original patient tumor.

### In vivo drug testing

*In vivo* drug studies were performed according to animal ethics approvals at TRI and QUT (QUT/275/17 and 1700000755). Mice were implanted with PDX fragments from 6-10 weeks of age. Once tumors reached ∼150-350 mm^3^ (faster models started drug between 150-250 mm^3^ and slower models between 250-350 mm^3^), mice were randomized into treatment groups and treated for 28 days via oral gavage. Efforts were made to have a similar number of mice on each arm from each passage carrying similar sized tumors. Mice were drugged 6 days on/1 day off with vehicle (20% Tween 20, 20% DMSO) or talazoparib (0.33mg/kg) as previously reported (17).

### DNA extraction and quality control

DNA was extracted from patient blood samples as well as patient and related PDX tumor samples, using DNeasy Blood & Tissue Kits (Qiagen, Germantown, MD, USA). The purity of DNA was assessed using NanoDrop and quantified using the Qubit dsDNA BR assay (Thermo Fisher Scientific, MA, USA). DNA samples were assayed with the Omni 2.5-8, V1.0 and V1.1 Illumina BeadChip as per manufacturer’s instructions (Illumina, San Diego, CA, USA). SNP array analysis to confirm sample identity, tumor content of DNA samples (18) for subsequent sequencing is described in detail in the Additional File 1:Supplementary Material.

### Whole exome and whole genome sequencing

Samples underwent whole-exome sequencing (WES) and whole-genome sequencing (WGS). The WES libraries were prepared using the SureSelect capture V5+UTR kit (Agilent, Santa Clara, CA, USA) and sequenced with 100bp paired-end sequencing on a HiSeq 2500/4000 (Illumina) to a targeted 100-fold read depth. The WGS libraries were prepared using the TruSeq Nano kit (Illumina) and sequenced with 150bp paired-end sequencing on a HiSeq X Ten (Illumina) at Macrogen (Geumcheon-gu, Seoul, South Korea) with targeted mean read depth of 60x for primary tumor samples and 30x for matched PDX and normal samples.

### Sequencing data analysis

Cutadapt (v1.18)(19) was used to trim low-quality 3’ bases (‘-q 20’) and remove adapters before alignment to a combined human/mouse (GRCh37/GRCm38 Nodshiltj background) reference using BWA-mem (v0.7.15)(20), and sorted and indexed using SAMtools (v1.9) (21). Duplicate reads were marked using Picard MarkDuplicates (v1.97). Human mutation calling process only used read-pairs aligned to the human sequences with a mapping quality score of 60. Quality assessment and coverage estimation was carried out by in-house developed tools, qProfiler and qCoverage. Downstream analysis included variant calling, copy number alteration (CNA) and structural variant (SV) detection (22-24), heterogeneity analysis (25, 26), microsatellite instability (MSI) and HR deficiency (HRD) status assessment (27, 28), and signature analysis (29, 30), and is described in detail in the Additional File 1:Supplementary Material.

Publicly available datasets (TCGA uterine corpus endometrial carcinoma (UCEC) and TCGA-UCS) were processed using a similar approach, without the initial human/mouse alignment and filtering step.

### Statistical analysis and data visualization

Statistical analysis and data visualization was performed in R 3.5.1, using ggplot2 and ComplexHeatmap packages, and using Circos. Final figure formatting was done with Illustrator (Adobe).

## Results

### Established PDX models represent four histological subtypes

Of 33 EC tumors implanted fresh, we generated 15 EC PDXs which were confirmed as EC models by a specialized anatomical pathologist. Successful engraftment rates were only obtained for histological grades 2 and 3 tumors implanted fresh (33 and 68%, respectively), none of grade 1 tumors engrafted (0/8) (Tables S1-2). We were also able to obtain 3 EC PDX models from an additional 20 EC tumors after storage at −80°C or 4°C overnight (Additional File 1:Table S2). In addition to 18 EC PDXs, seven models showed *in vivo* tumor growth, however these were confirmed to be lymphomas based on positive leukocyte common antigen staining. This occurred more often from grade 1 tumors (3/14, 21%) than grade 3 ECs (3/28, 11%). This study reports detailed genomics data for 11 of 18 EC PDX models.

The 11 PDX models were from EC of patients with a mean age of 70 (range 43-86 years, Additional File 1:Table S2) who represented the wide range of EC disease with varying histology and stages (IA to IIIB). Histologic diagnoses included carcinosarcoma (n=3), mixed endometrioid and serous (n=2), mixed endometrioid and clear cell (n=1) and endometrioid (n=5, of which 4 were FIGO grade 3 and 1 FIGO grade 2). Nine patients received radiation or chemoradiation and six patients recurred, five of which have subsequently died. In all models, the tissue architecture, the epithelial compartment and the global histological classification features were preserved in the corresponding F0 to F2 PDXs (Fig. 1).

**Fig. 1.**
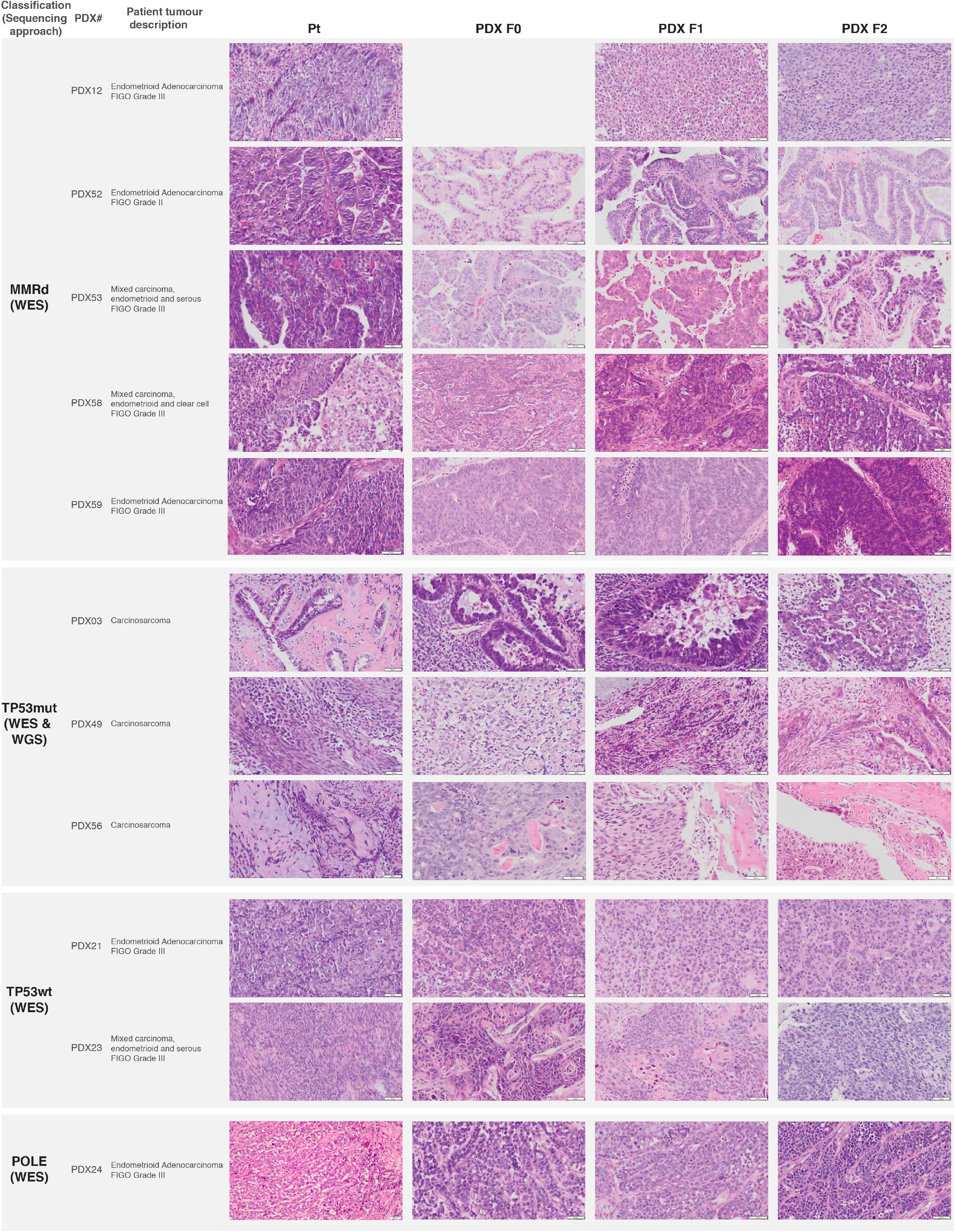
Histopathology assessment of the primary and PDX tumor samples. PR: primary tumor. PDX-F0: 1^st^ tumor obtained from the mice transplant. PDX-F1, PDX-F2: subsequent transplants. F0 picture for PDX12 is missing as no FFPE sample was available for this lineage.

### Genomics of EC PDX models

Sequencing and SNP array analysis were performed to molecularly classify the 11 EC PDX models. The molecular classification was adapted from the TCGA endometrial and UCS studies, and was based on five aspects (Fig. 2): commonly mutated genes (Additional File 1:Table S3, Additional File 2: Table S4), TMB, MSI score (Figure S1), extent of genomic CNA (Additional File 1:Fig. S2), and mutational signatures (Additional File 1:Fig. S3). Four molecular subtypes were represented in the generated PDX models (Fig. 2, Additional File 1:Fig. S4). One PDX model was *POLE*-mutated. It contained p.Pro286Arg *POLE* mutation in the exonuclease domain, previously reported in EC and shown to lead to a particularly strong mutator phenotype (31). This PDX was characterized by an ultra-high TMB (>600 Mutations/Mb), a CNA stable genome, low MSI score (<3.5%) and a *POLE*-associated mutational signature. Two PDX models were classified as *TP53* wild-type (p53wt), since they lacked *TP53* or *POLE* mutations and were MSI-stable. They were characterized by a relatively low TMB, and moderately stable genomes. Five PDX models were classified as MMRd, based on a high TMB (>20 Mutations/Mb), high MSI scores, and MMRd-associated mutational signatures.

**Fig. 2.**
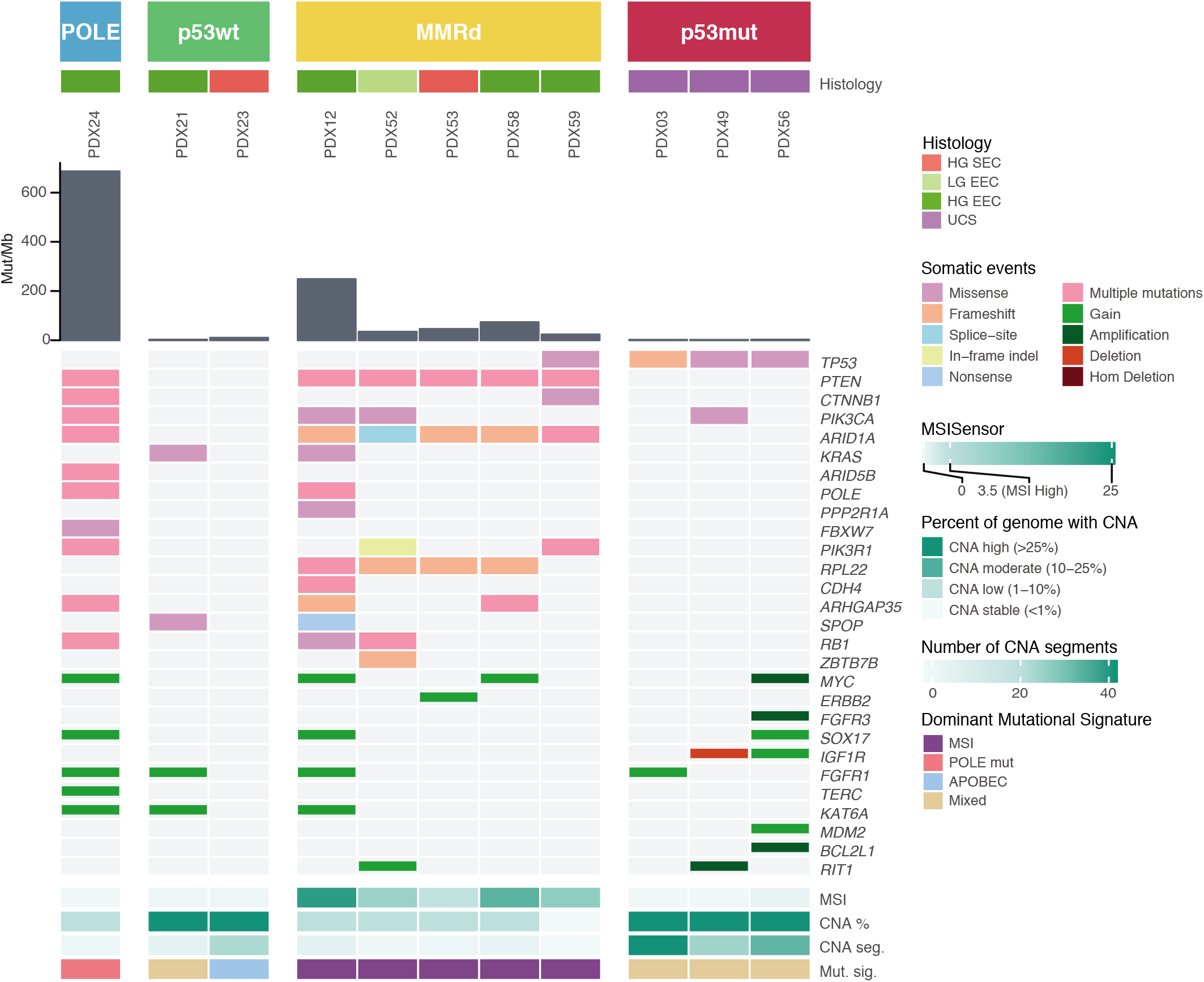
The four molecular subtypes are represented in PDX models. Genomic characteristics of endometrial carcinoma and carcinosarcoma PDX models. PDX models are grouped by the four molecular subtypes: POLE, p53wt, MMRd and p53mut. Tumour mutation burden is shown by grey bars, as mutations per Mb. Somatic mutations and CNA events, which were detected in PDX samples in genes relevant to endometrial carcinomas and carcinosarcomas (Additional File 1:Table S3, Additional File 2:Table S4), are shown. Only consensus variants detected in all sequenced PDX tumor samples were included in this figure. MSI score was assessed by MSISensor (Additional File 1:Fig. S1). Percentage of genome with CNA and the number of CNA segments were determined from SNP arrays or WGS data (Additional File 1:Fig. S2). Only the dominant mutational signature etiology is shown.

All three UCS models were classified as *TP53* mutant (p53mut), as they all harbored hotspot or deleterious somatic *TP53* mutations, low TMB (<10 Mutations/Mb), high degree of genomic instability (>25% of genome with CNAs and >15 CN segments) and low MSI scores, suggesting microsatellite stability. The UCS models had a mixed mutational signature profile, with no dominant signature detected (<30% of somatic mutations attributed to a single signature; Additional File 1:Fig. S3).

Molecular subtyping of two models with serous histology revealed one was p53wt tumor (PDX23) and another was MMRd (PDX53), which was consistent with a germline *MSH6* mutation (p.Tyr214*) in the latter patient and the finding that 13% of patients with germline MMR mutations have a mixed serous histology (32). Somatic mutations detected in other genes were consistent with TCGA findings. Protein altering somatic variants were commonly detected in *PTEN, ARID1A, PIK3CA, KRAS* and *CTNNB1* genes. All MMRd models contained somatic *PTEN* and *ARID1A* mutations, most of which were inactivating frameshift or nonsense mutations (Fig. 2, Additional File 2:Table S4).

### Variable intra-tumor heterogeneity observed in MMRd PDX models

To study the intra-tumor heterogeneity and to evaluate how well the PDX models recapitulate primary tumors, we focused in detail on four MMRd models as these might be expected to accumulate changes during passaging based on their defective DNA mismatch repair. Genome-wide levels of CNA and LOH changes were comparable between primary and PDX tumor samples (Fig. 3a, Additional File 1:Fig. S5). The TMB between primary tumor and different passages of PDX (passage 0-4) was stable across different passages of PDX samples; however we observed a substantially higher number of mutations in PDX samples compared with matched primary samples in three PDX models (Fig. 3b). This was likely due to lower tumor purity observed in the primary samples compared with PDX samples (Additional File 1:Fig. S6). Indeed, PDX59 with the highest tumor purity had the most comparable number of somatic mutations to the matched PDX samples. For three of four models, 83-99% of the somatic substitutions in the primary tumor were also detected in all tested PDX samples, with only limited heterogeneity observed between different lineages of the PDX (Fig. 3c and Additional File 1:Fig. S7). In PDX58 however, we observed that the established PDX shared only a third of its somatic substitutions with the primary tumor sample (Fig. 3d). A clonality analysis of PDX58 using PyClone identified two distinct mutational clones (PDX lineages A and B) that likely diverged early in the tumor evolution and were unintentionally selected during the initial tumor transplantation (Fig. 3e-f). Mutations unique to lineage A included an activating *KRAS* mutation (p.Gly12Asp) and a hotspot *TP53* mutation (p.Arg273His). Overall, since the greatest variability was observed between different lineages of the established PDX models and not between the passages, we concluded that this was due to spatial heterogeneity present in the original patient tumor.

**Fig. 3.**
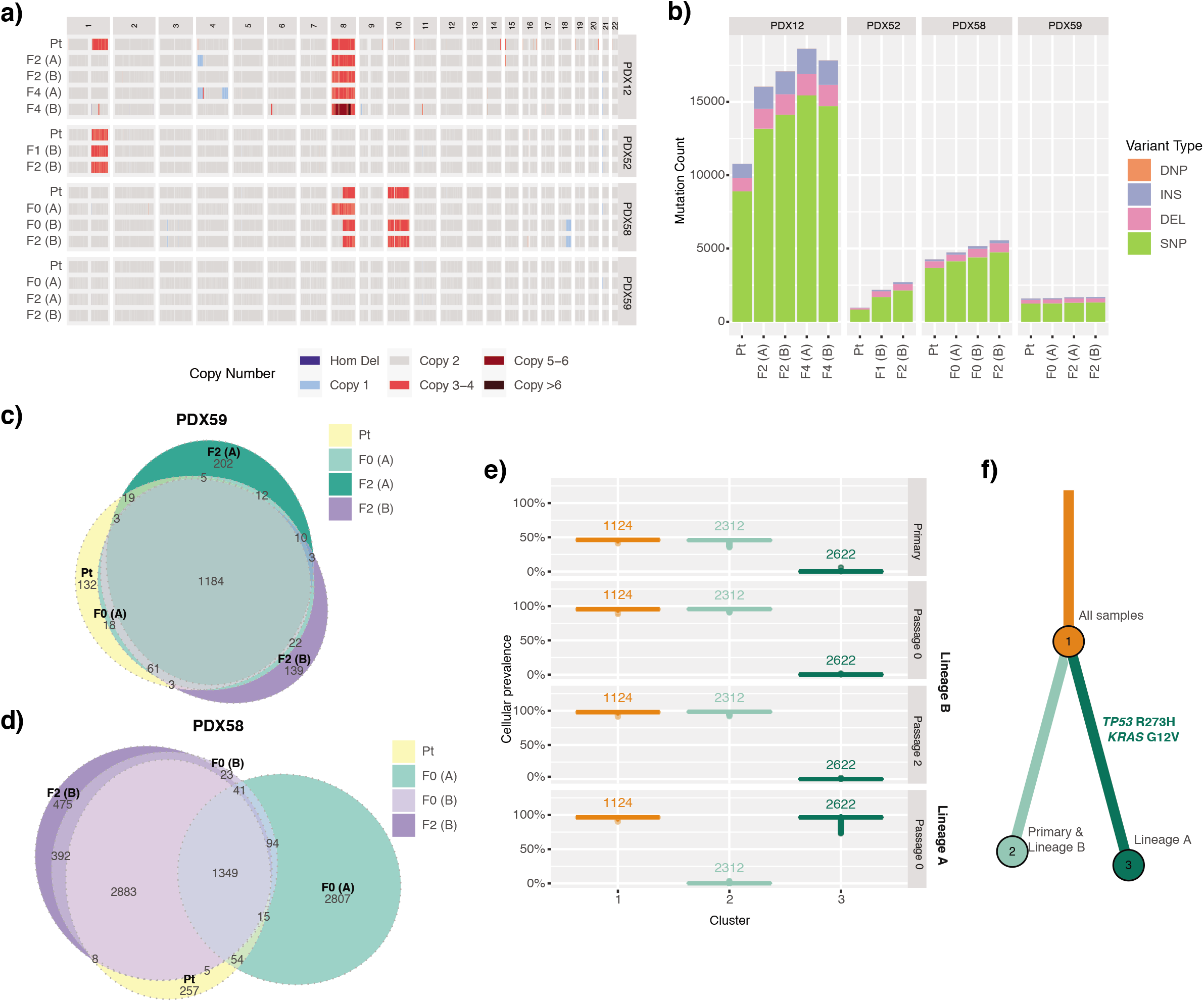
Intra-tumor heterogeneity observed in the MMRd EC PDX models. **a** Genome-wide levels of CNA and **b** total somatic mutation count in the four MMRd models, where primary tumor sample was analyzed by WES and by SNP arrays. Varying degrees of mutational heterogeneity visualized by Euler diagrams of somatic substitutions called by qBasepileup in **c** PDX59 and **d** PDX58 MMRd models. **e** Cellular prevalence and **f** the clonal evolution tree of the top three mutational clusters (with ≥5% of all somatic substitutions) detected in the PDX58 model by PyClone. Values shown above boxplots represent the number of substitutions contributing to each cluster. Length of branches is proportional to the number of substitutions attributed to that clone. Tumor samples are grouped by patient ID. PDX samples are labelled by passage number and lineage in brackets. DEL — deletion; DNP — double nucleotide polymorphism; INS — insertion; SNP — single nucleotide polymorphism; Hom Del — homozygous deletion.

### Variable intra-tumor heterogeneity observed in p53mut uterine carcinosarcoma

*PDX models* To capture the genome instability and heterogeneity observed in the *p53*mut UCS models, we performed WGS to examine CNA changes in more detail. The overall CNA and LOH changes were comparable between the primary and matched PDX samples, with the exception of PDX03 model (Fig. 4a, Additional File 1:Fig. S8), which harbored a whole-genome duplication (WGD) not detected in the primary tumor. The total number of somatic mutations detected across the whole genome for each UCS model was consistent between the primary tumor and the matched PDX samples, with no increase in mutation number detected with passaging (Fig. 4b).

**Fig. 4.**
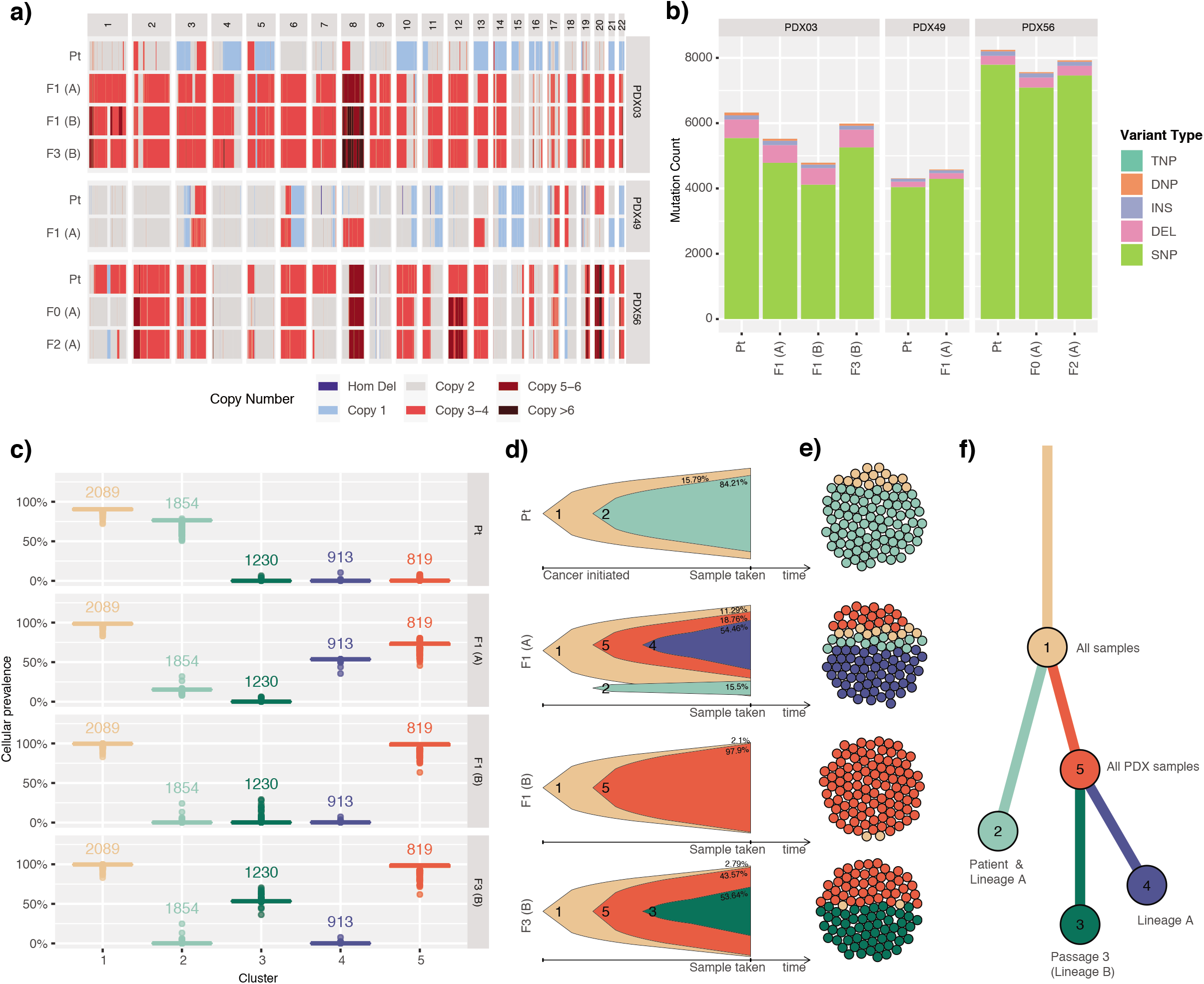
Intra-tumor heterogeneity and clonal evolution observed in p53mut UCS PDX models. **a** Genome-wide levels of CNA and **b** total somatic mutation count in the three UCS models. **c** Cellular prevalence of the top five mutational clusters with ≥5% of all somatic substitutions detected in the PDX03 model by PyClone. Values shown above boxplots represent the number of substitutions contributing to each cluster. **d** Fish plots and **e** cellular population depictions of the top five mutational clusters detected in the PDX03 carcinosarcoma model. Percentages shown in the fish plots are the estimated proportions of cells containing that mutational cluster. **f** The clonal evolution tree inferred by ClonEvol, where length of branches is proportional to the number of substitutions attributed to that clone. TNP — triple nucleotide polymorphism.

For PDX03, which contained a WGD in the PDX and not the matched primary tumor, PyClone clonality analysis revealed a high degree of heterogeneity in the model, with five major different mutational clones detected (Fig. 4c-e) that were associated with multiple samples in PDX lineages A and B. Clone 2 was the predominant clone in the primary tumor sample (predicted in around 80% of tumor cells), but was detected at only around 15% in lineage A, and was absent in lineage B. Interestingly, using Battenberg we identified a CNA subclone in lineage A with a similar copy number profile to the primary tumor (Additional File 1:Fig. S9a-d). In support of this, the ploidy estimated from SNP array analysis of the additional PDX samples, identified two early lineage samples (passage 0 and 1) from lineage A that had estimated ploidy of 2, same as the primary tumor sample (Additional File 1:Fig. S9e). We therefore concluded that the WGD event was already present in a subclone of the primary tumor, although not detected in the sample taken for WGS analysis. In the other two carcinosarcoma models, clonality analysis also revealed heterogeneity, although not to the same extent as seen in the PDX03 model (Additional File 1:Fig. S10-11).

### Assessment of PARPi responses and HR status in PDX models

Since the established PDX models were found to reflect the primary tumors, we evaluated their use for testing molecularly-targeted treatments. Potential therapeutic options for all 11 PDX models were identified using their genomic profiles and Cancer Genome Interpreter analysis (Additional File 1:Fig. S12). Multiple therapeutic options were identified, with three or more options detected per PDX model. PARP, mTOR and PI3K pathway inhibitors, as well as PD1 inhibitors were identified among the most common potential treatment options. However, a large number of therapeutic agents were identified for tumors with a high TMB (POLE/MMRd), where most somatic variants could be passenger events, thus functional testing is required to determine whether these models are responsive.

PARPi has been previously identified as a promising therapeutic strategy for EC. Therefore, we evaluated PARPi responses in a subset of the established PDX models. Highly potent PARPi talazoparib was selected due to its significantly higher PARP trapping ability (33) and was used to treat three UCS PDX models, one p53wt model and two MSI models *in vivo* (Fig. 5). PDX03 and PDX49 showed the most sensitivity, with average tumor growth inhibition (TGI) showing the equivalent of stable disease by Response Evaluation Criteria In Solid Tumors (RECIST) criteria. PDX56 and PDX23 showed significant TGI, albeit this would translate to progressive disease by RECIST criteria. No effect of talazoparib was seen in the MMRd models with multiple heterozygous missense mutations and/or LOH in DNA repair genes (PDX12 and PDX53).

**Fig. 5.**
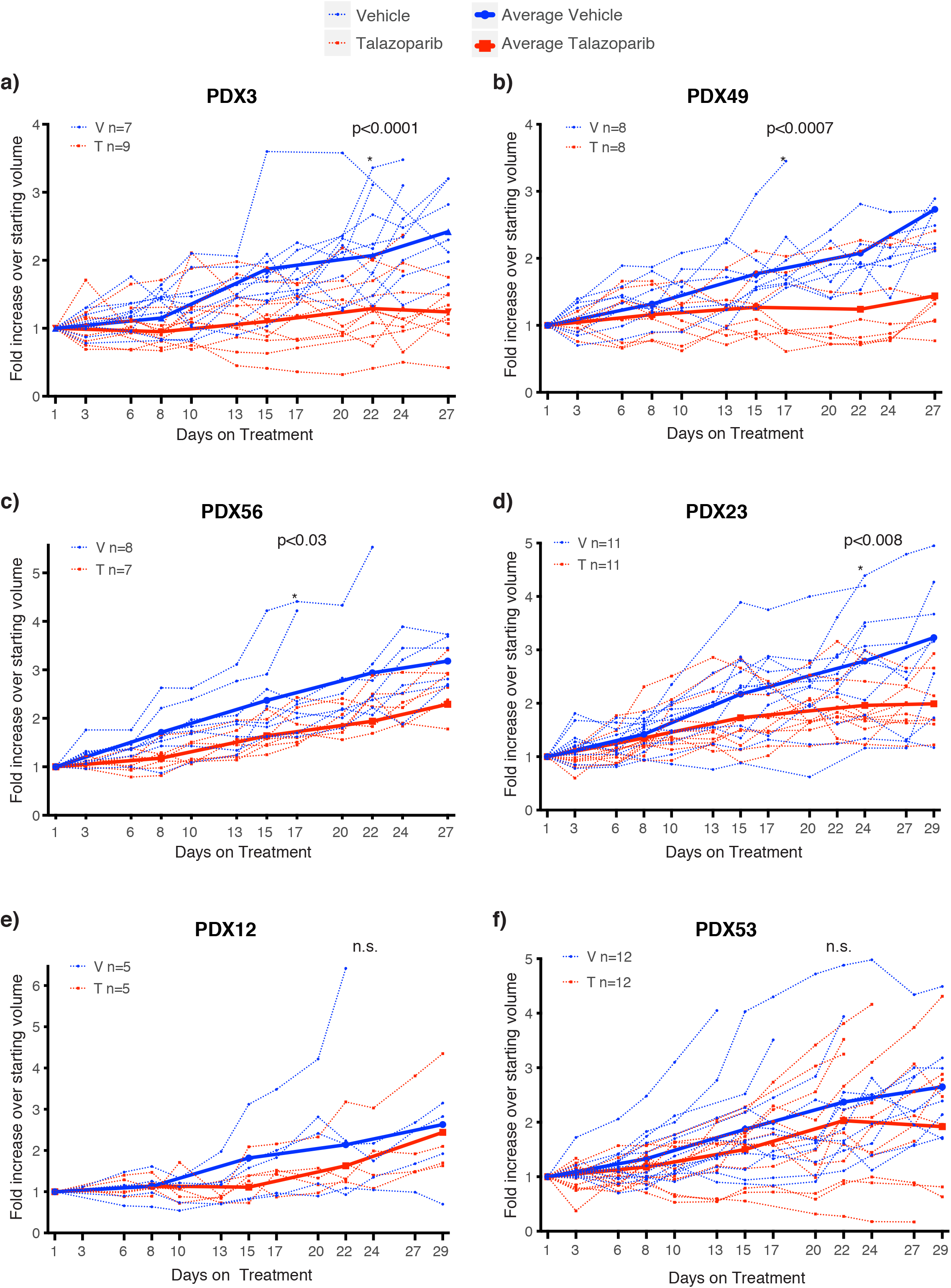
Talazoparib responses in EC and UCS PDX models. Talazoparib responses in **a** PDX03 — p53mut UCS; **b** PDX49 — p53mut UCS; **c** PDX56 — p53 UCS with somatic *ARID1A* deletion; **d** PDX23 — p53wt EC; **e** PDX12 — MMRd EC with somatic *PTEN, BRCA2, ATM* and *PALB2* mutations; **f** PDX53 — MMRd EC with somatic *PTEN, ATM, BRCA1* and *MRE11A* mutations. Recipient mice bearing PDX at starting volume of ∼150-350 mm^3^ were randomized to treatment with vehicle or talazoparib (0.33mg/kg) for 28 days (6 days on, one day off) via oral gavage. Analysis for significance between treatment groups was performed using a repeated mixed effects analysis (which can account for random missing measurements) on the day the first mouse was sacrificed based on tumor size (e.g. 17, 22 and 24 days), except for PDX53 where 2 vehicle mice were sacrificed early and excluded. n.s — not significant; * — significant difference (p-value shown).

Due to the limited responses to single agent PARPi observed *in vivo*, we characterized the PDX HR status in detail. We firstly estimated the HRD scores for the primary and matched PDX samples with WGS or SNP array-estimated CNA data. All of the primary tumor samples had HRD scored below the threshold of ≥42, a previously defined cut-off for HRD breast and ovarian cancers (34, 35). Two of three carcinosarcoma p53mut models and one p53wt model had some PDX samples with HRD scores just over the threshold (Fig. 6a).

**Fig. 6.**
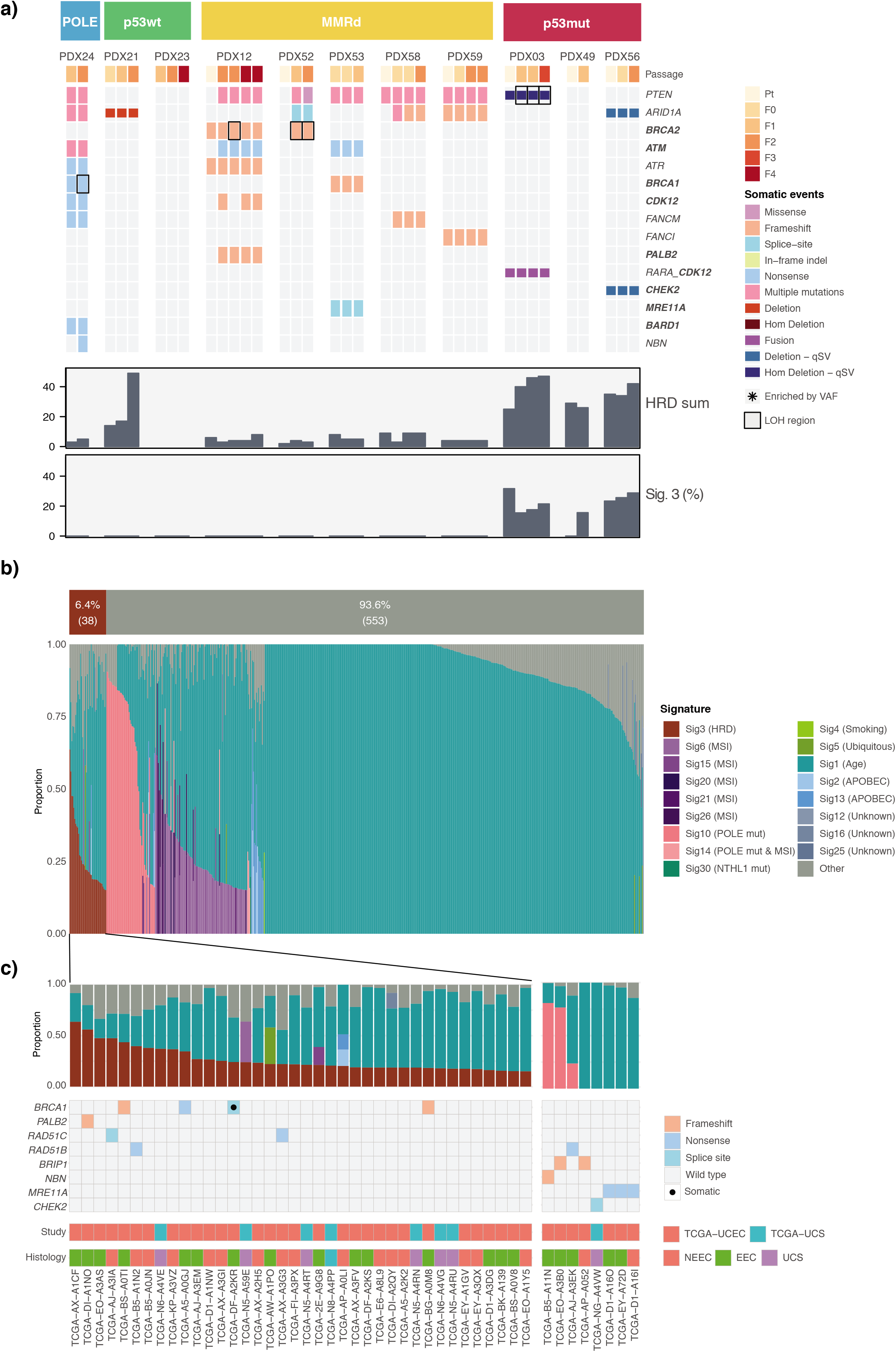
Genomic HRD assessment in EC PDX models and public data. **a** HRD assessment in PDX models. Somatic substitutions, indels, CNAs and SVs are shown for DNA repair related genes, including *PTEN* and *ARID1A* (Additional File 1:Table S5, Additional File 2:Table S6). HR-related genes are highlighted in bold. No pathogenic or likely pathogenic germline substitution and indel variants in these genes were detected. HRD sum scores were determined using scarHRD from SNP arrays and WGS data, where available. Percentage of Signature 3 was determined with deconstructSigs using COSMIC v2 signatures. Only WGS data is shown for PDX03 and PDX49, where WES and WGS was performed. **b** Mutational signature assignment for TCGA-UCEC and TCGA-UCS cohorts (n=591). Signature assignment was performed using deconstructSigs with 15% minimum signature cut-off. **c** TCGA-UCEC and TCGA-UCS cases with possible HRD. Cases with pathogenic or likely pathogenic variants in HR-related genes (Additional File 1:Table S5, Additional File 2:Table S7) or cases with Signature 3 detected are included.

We then assessed the mutational signatures, specifically contribution of COSMIC Signature 3, as an alternative marker of HRD. All three carcinosarcoma models had a low mutational contribution of Signature 3, between 20-30% (Fig. 6a). However, rearrangement signature analysis on the three carcinosarcoma models, did not detect *BRCA1/2-*associated signatures (Additional File 1:Fig. S13). Finally, we looked for the presence of germline and somatic mutations in the HR-associated genes (Additional File 1:Table S5) and somatic mutations in *PTEN* and *ARID1A*, since mutations in these genes have been associated with PARPi sensitivity in the pre-clinical setting (11, 14). Apart from recurrent damaging mutations in *PTEN* and *ARID1A*, we did not detect any clear driver mutations in HR-genes with evidence of enrichment or LOH in p53mut or p53wt groups. We did detect a large number of likely-passenger mutations in hypermutated models (MMRd/ POLE; Fig. 6a; Additional File 2:Table S6). The genomic characterization and the lack of *in vivo* tumor regressions in response to PARPi suggested that none of the PDX models had clear evidence of HRD.

### HR deficiency is predicted to be a rare event in UCEC and UCS

Since we did not observe clear evidence of HRD in the PDX models, we wanted to estimate the rate of HRD in a larger cohort of unselected UCEC and UCS patients. We assessed HR status in the TCGA studies of endometrial carcinomas (TCGA-UCEC; n=560) and uterine carcinosarcomas (TCGA-UCS; n=57). *De novo* detection of mutational signatures identified six signatures associated with age, APOBEC, MMR and POLE, but not HRD-associated Signature 3 in either of the datasets (Additional File 1:Fig. S14). We then applied the same approach that was used for the PDX models — assigning known COSMIC signatures (including HRD Signature 3) to the mutational profiles of each sample. Only samples with >50 somatic substitutions were selected for this analysis (n=591). Using this approach, we predicted that only 6.4% (38 of 591) of all analyzed samples had Signature 3 detected, using 15% minimum signature contribution cut-off to avoid overfitting (Fig. 6b). Signature 3 was detected in 11.4% (16/141) of non-endometrioid EC, 3.8% (15/393) of endometrioid EC and 12.3% (7/57) of UCS cases.

We also looked for germline and somatic variants in HR genes in these 591 samples (Fig. 6c, Additional File 2:Table S7). Only 2.5% (15/591) cases were found to harbor germline pathogenic or likely pathogenic HR gene variants: 3.5% (5/141) of non-endometrioid, 2.3% (9/393) of endometrioid and 1.8% (1/57) of UCS cases. Seven variants in *BRCA1, PALB2, RAD51C* and *RAD51D* genes were concurrent with the presence of mutational Signature 3; whereas the variants in *BRIP1, NBN, MRE11A* and *CHEK2*, as well as one variant in *RAD51B*, were detected in cases without Signature 3. This finding was in line with a previous report in breast cancer, where mutations in DNA-damage signaling pathway genes, such as *ATM* and *CHEK2*, were not associated with increased Signature 3 (36). Somatic mutation analysis was restricted to 38 cases with detected Signature 3. Only one somatic pathogenic variant in *BRCA1* was detected in endometrioid cancer.

## Discussion

To understand the suitability of pre-clinical PDXs models to study EC, we undertook an in-depth genomic characterization of patient primary and matched serial PDX tumor samples of EC. Although some molecular typing of EC PDXs has been previously reported (e.g. those with microsatellite stability versus instability) (37, 38), this is the first report of *de novo* mutational signature and copy number analysis across a panel of EC PDXs. The established PDX models recapitulated key morphological and genomic features present in the molecular subtypes (3). Interestingly, no distinct PDX mutational signatures were found using *de novo* signature analysis, and the mutational profiles were very similar between the primary and matched PDX samples. We also did not observe an accumulation of PDX-specific CNA events, as was previously reported in PDX models of breast, brain, lung, colon and pancreatic cancers(39). Taken together, these results support that PDX-specific tumor evolution is minimal in these models, and they reliably represent the primary tumors. PDX engraftment was much higher in G3 tumors. For future studies, the addition of one dose of rituximab during implantation could reduce the incidence of lymphomas as previously reported in ovarian cancer(40).

### Intra-tumor heterogeneity

EC tumors are composed of multiple complex sub-clonal cell populations resulting in intra-tumor heterogeneity(41, 42). Maximum tolerated dose chemotherapy regimens aim to eradicate the entire tumor but rarely achieve it, often leaving resistant sub-clones that possess a growth advantage and are free to expand. Hence, genomic intra-tumor heterogeneity has clinical implications for EC, and needs to be characterized to enable precision medicine. In this study we transplanted undisturbed tumor fragments and characterized multiple PDX lineages, which allowed us to observe great variation in the intra-tumor heterogeneity among different PDX models.

In two of four models where multiple lineages were sequenced we detected high levels of intra-tumor heterogeneity that could have a potential impact on treatment responses. The greatest variability was observed between the different lineages (detected in early passages) and not between the passages, indicating that this heterogeneity was pre-existing in the primary tumor, although it is unclear whether the subclones were selected due to chance or a selective advantage in the PDX. In the MMR-deficient model (PDX58), one of the established lineages was enriched for a subclone with hotspot *KRAS* and *TP53* mutations and over 50% private mutations, suggesting that this subclone had diverged early on in the tumor evolution. *KRAS* mutations have been linked to drug resistance in multiple cancers (43, 44), and *TP53* mutations are associated with poor prognosis in EC (3). In the *TP53*-mutant UCS model (PDX03), one of the lineages had a WGD event together with other subclonal mutations. WGDs are frequently detected in UCS (90%) (10) compared with epithelial EC, as well as in metastatic cancers across multiple cancer types (45) and have been associated with poor prognosis (46). PDX models that capture pre-existing intra-tumor heterogeneity such as described here, make a perfect tool for studying the effects of individual genomic events on tumor evolution, progression and drug responses, and should be explored further.

### PARP inhibitors in EC

PARPi sensitivity has previously been reported in EC, although to date the work has been performed in cell lines, which do not faithfully represent all of the EC molecular subtypes (11, 13). The proposed biomarkers of PARPi response in EC are diverse, including *PTEN, ARID1A* or *MRE11A* loss (11, 13, 14), *TP53* mutations, and cumulative effect of multiple somatic hits in HR genes in hypermutated MMRd EC. Our PDX EC cohort had a representation of all of these events, so we could investigate their effect on PARPi response *in vivo*. By performing HRD scarring and mutational signature assessment, commonly used for classifying HRD-ness in ovarian and breast cancers (47), we determined that our EC models were all likely HR-proficient. Interestingly, three UCS models with mutated *TP53* had intermediate HRD scores and some of the somatic mutations were attributed to Signature 3, although it was not the dominant signature. Two of these UCS models also showed disease stabilization in response to the potent PARPi talazoparib *in vivo*. The PDX models of other molecular subtypes did not have a marked response to talazoparib (p53wt and MMRd with multiple damaging mutations in canonical HR genes, although none with consistent enrichment in tumors). Furthermore, mutations in *PTEN* or *ARID1A* did not have an effect on PARPi response in our models.

It has been recently reported that up to half of non-endometrioid EC (predominantly p53mut) can harbor BRCA-associated genomic scars compared with only 12% of endometrioid ECs (p53wt/MMRd) (16). Since our PDX cohort did not have a large representation of non-endometrioid EC (five of 11 models), it was possible that we missed the HRD cases by chance. However, our exploration of TCGA UCEC and UCS datasets also showed that HRD is likely rare in EC. We saw much lower rates of Signature 3 both in general (6.8%), and in the non-endometrioid cancers (12%) compared to the previous report (16). This was likely due to a more conservative signature assignment approach used in our study. The mutational signature analysis approach can have a great influence on the identification of signatures, especially signatures with flat profiles (Signature 3, 5 and 8) (48). Furthermore, the rates of damaging mutations in HR genes were quite rare (1.6-3.6% depending on the histological subtype), consistent with another study looking at bi-allelic alterations in HR genes (15). The lack of tumor regressions in our EC PDX models in response to PARPi talazoparib and the infrequent HRD events in EC public datasets indicate that PARPi may not be sufficient as a single agent therapy in an unselected EC patient population. Nonetheless, PARPi may still have an important role to play in the management of EC, and should be further investigated in combination with other treatments. Several PARPis including talazoparib have been shown to have strong PARP trapping effect (33), leading to replication stalling. This opens up the possibility to combine PARPi with other therapies for an enhanced anti-tumor activity, including cell cycle checkpoint inhibition, RNA Pol1 inhibition (49) or ICIs. Genomically-characterized PDX models, such as ours, will be crucial for assessing the efficacy of these combinations in EC, as has already been assessed in other cancers (49, 50). Importantly though, PARPi and ICI combinations will need to be assessed in humanized PDX models, as regular PDX models lack representation of the immune landscape.

## Conclusions

In conclusion, we have shown that EC PDX models can capture intra-tumor heterogeneity, which should be accounted for and explored to improve treatment responses and patient outcomes. By combining genomic characterization and *in vivo* treatments, we also showed that PARPi talazoparib had disease stabilization activity in *TP53*-mutant EC, which can potentially be enhanced by combination therapies.

## List of Abbreviations

CAN: Copy number alteration
EC: Endometrial cancer
FFPE: Formalin-fixed paraffin-embedded
H&E: Haematoxylin and eosin
HR: Homologous recombination
HRD: HR deficiency
ICI: Immune checkpoint inhibitor
LOH: Loss of heterozygosity
MMRd: Mismatch repair deficient
MSI: Microsatellite instability
NSG: Nod scid gamma
p53mut: TP53 mutant
p53wt: TP53 wildtype
PARPi: PARP inhibitor
PDX: patient-derived xenograft
RECIST: Response Evaluation Criteria In Solid Tumors
SV: Structural variant
TCGA: The Cancer Genome Atlas
TGI: tumor growth inhibition
TMB: Tumor mutation burden
UCEC: Uterine corpus endometrial carcinoma
UCS: Uterine carcinosarcoma
WES: Whole-exome sequencing
WGD: whole-genome duplication
WGS: Whole-genome sequencing

## Declarations

### Ethics approval and consent to participate

All samples were obtained from patients with informed consent and the study has human ethics approval from the Mater Health Services Human Research Ethics Committee (HREC/15/MHS/127), UnitingCare Health Human Research Ethics Committee (1116), Queensland University of Technology (1500000169, 1500000323) and QIMR Berghofer (P3478, P2095).

PDX establishment and passaging was performed according to animal ethics approvals at TRI (TRI/021/19) and QUT (1900000701). In vivo drug studies were performed according to animal ethics approvals at TRI and QUT (QUT/275/17 and 1700000755).

## Consent for publication

Not applicable.

## Availability of data and materials

The datasets (raw data files for WGS and WES) supporting the conclusions of this article are available in the European-Genome Phenome Archive under study number EGAS00001004666. Access to SNP array data can be requested by contacting the corresponding author.

The code required to reproduce the figures in this manuscript can be found on https://github.com/okon/EC_PDX_genomics. The code used to perform alignment to human and mouse genome references is available on https://github.com/ampatchlab/nf-pdx. The in-house tools are available on https://github.com/AdamaJava.

## Authors’ contributions

PP designed and supervised the study, provided study resources, managed the project, performed experiments and data analysis, and wrote the manuscript. NW and A-MP designed and supervised the study, managed the project and wrote the manuscript. VFB designed the study, developed the methodology, performed *in vivo* experiments and corresponding data analysis, and wrote the manuscript. OK developed the methodology, supervised and performed genomic data analysis, and wrote the manuscript. KN, LTK, SHK, ALD, PR-M, VL and FN performed genomic data analysis. ATS performed data curation, experiments and data analysis. LMP provided study resources, performed experiments and data analysis. RJ performed experiments and data analysis. DS performed histopathology data analysis. RR and CD provided study resources, curated data and managed the project administration. JN, AG, NC and LP provided study resources and curated data. JVP provided study resources and developed the methodology. All authors reviewed and approved the final manuscript.

## Acknowledgements

Authors would like to thank all patients included in this study. We are also grateful for the staff at the Translational Research Institute of Australia (TRI) Biological Research Facility and TRI microscopy facility. The Translational Research Institute is supported by a grant (APP108382) from the Australian Government. We would also like to acknowledge members of the Medical Genomics and Genome Informatics teams at the QIMR Berghofer Medical Research for their technical support. The results shown here are in part based upon data generated by the TCGA Research Network: https://www.cancer.gov/tcga.

## Additional files

**Additional File 1:** pdf; Supplementary Methods, Supplementary Tables 1-3,5 and Supplementary Figures.

**Additional File 2:** xlsx; Supplementary Tables 4,6-8 (descriptions provided in the file).

## References

1. Siegel RL, Miller KD, Jemal A. Cancer statistics, 2018. CA: a cancer journal for clinicians. 2018;68(1):7–30.

2. Stelloo E, Nout RA, Osse EM, Jurgenliemk-Schulz IJ, Jobsen JJ, Lutgens LC, et al. Improved Risk Assessment by Integrating Molecular and Clinicopathological Factors in Early-stage Endometrial Cancer-Combined Analysis of the PORTEC Cohorts. Clin Cancer Res. 2016;22(16):4215–24.

3. Levine DA, Network CGAR. Integrated genomic characterization of endometrial carcinoma. Nature. 2013;497(7447):67–73.

4. Talhouk A, McConechy M, Leung S, Li-Chang H, Kwon J, Melnyk N, et al. A clinically applicable molecular-based classification for endometrial cancers. British journal of cancer. 2015;113(2):299–310.

5. Talhouk A, McConechy MK, Leung S, Yang W, Lum A, Senz J, et al. Confirmation of ProMisE: a simple, genomics-based clinical classifier for endometrial cancer. Cancer. 2017;123(5):802–13.

6. Kommoss S, McConechy M, Kommoss F, Leung S, Bunz A, Magrill J, et al. Final validation of the ProMisE molecular classifier for endometrial carcinoma in a large population-based case series. Annals of Oncology. 2018;29(5):1180–8.

7. Cosgrove CM, Tritchler DL, Cohn DE, Mutch DG, Rush CM, Lankes HA, et al. An NRG Oncology/GOG study of molecular classification for risk prediction in endometrioid endometrial cancer. Gynecologic oncology. 2018;148(1):174–80.

8. Jones S, Stransky N, McCord CL, Cerami E, Lagowski J, Kelly D, et al. Genomic analyses of gynaecologic carcinosarcomas reveal frequent mutations in chromatin remodelling genes. Nature communications. 2014;5(1):1–7.

9. Zhao S, Bellone S, Lopez S, Thakral D, Schwab C, English DP, et al. Mutational landscape of uterine and ovarian carcinosarcomas implicates histone genes in epithelial– mesenchymal transition. Proceedings of the National Academy of Sciences. 2016;113(43):12238–43.

10. Cherniack AD, Shen H, Walter V, Stewart C, Murray BA, Bowlby R, et al. Integrated molecular characterization of uterine carcinosarcoma. Cancer cell. 2017;31(3):411–23.

11. Dedes KJ, Wetterskog D, Mendes-Pereira AM, Natrajan R, Lambros MB, Geyer FC, et al. PTEN deficiency in endometrioid endometrial adenocarcinomas predicts sensitivity to PARP inhibitors. Science translational medicine. 2010;2(53):53ra75–53ra75.

12. Miyasaka A, Oda K, Ikeda Y, Wada-Hiraike O, Kashiyama T, Enomoto A, et al. Anti-tumor activity of olaparib, a poly (ADP-ribose) polymerase (PARP) inhibitor, in cultured endometrial carcinoma cells. BMC cancer. 2014;14(1):179.

13. Koppensteiner R, Samartzis EP, Noske A, von Teichman A, Dedes I, Gwerder M, et al. Effect of MRE11 loss on PARP-inhibitor sensitivity in endometrial cancer in vitro. PloS one. 2014;9(6):e100041.

14. Shen J, Peng Y, Wei L, Zhang W, Yang L, Lan L, et al. ARID1A deficiency impairs the DNA damage checkpoint and sensitizes cells to PARP inhibitors. Cancer discovery. 2015;5(7):752–67.

15. Riaz N, Blecua P, Lim RS, Shen R, Higginson DS, Weinhold N, et al. Pan-cancer analysis of bi-allelic alterations in homologous recombination DNA repair genes. Nature communications. 2017;8(1):857.

16. de Jonge MM, Auguste A, van Wijk LM, Schouten PC, Meijers M, ter Haar NT, et al. Frequent homologous recombination deficiency in high-grade endometrial carcinomas. Clinical Cancer Research. 2019;25(3):1087–97.

17. Shen Y, Rehman FL, Feng Y, Boshuizen J, Bajrami I, Elliott R, et al. BMN 673, a novel and highly potent PARP1/2 inhibitor for the treatment of human cancers with DNA repair deficiency. Clinical Cancer Research. 2013;19(18):5003–15.

18. Song S, Nones K, Miller D, Harliwong I, Kassahn KS, Pinese M, et al. qpure: A tool to estimate tumor cellularity from genome-wide single-nucleotide polymorphism profiles. PloS one. 2012;7(9):e45835.

19. Martin M. Cutadapt removes adapter sequences from high-throughput sequencing reads. EMBnet journal. 2011;17(1):10–2.

20. Li H. Aligning sequence reads, clone sequences and assembly contigs with BWA-MEM. arXiv preprint arXiv:13033997. 2013.

21. Li H, Handsaker B, Wysoker A, Fennell T, Ruan J, Homer N, et al. The Sequence Alignment/Map format and SAMtools. Bioinformatics. 2009;25(16):2078–9.

22. Patch AM, Christie EL, Etemadmoghadam D, Garsed DW, George J, Fereday S, et al. Whole-genome characterization of chemoresistant ovarian cancer. Nature. 2015;521(7553):489–94.

23. Raine KM, Van Loo P, Wedge DC, Jones D, Menzies A, Butler AP, et al. ascatNgs: Identifying Somatically Acquired Copy-Number Alterations from Whole-Genome Sequencing Data. Current protocols in bioinformatics. 2016;56(1):15.9. 1-.9. 7.

24. Popova T, Manié E, Stoppa-Lyonnet D, Rigaill G, Barillot E, Stern MH. Genome Alteration Print (GAP): a tool to visualize and mine complex cancer genomic profiles obtained by SNP arrays. Genome biology. 2009;10(11):1–14.

25. Roth A, Khattra J, Yap D, Wan A, Laks E, Biele J, et al. PyClone: statistical inference of clonal population structure in cancer. Nature methods. 2014;11(4):396–8.

26. Dang H, White B, Foltz S, Miller C, Luo J, Fields R, et al. ClonEvol: clonal ordering and visualization in cancer sequencing. Annals of oncology. 2017;28(12):3076–82.

27. Niu B, Ye K, Zhang Q, Lu C, Xie M, McLellan MD, et al. MSIsensor: microsatellite instability detection using paired tumor-normal sequence data. Bioinformatics. 2014;30(7):1015–6.

28. Sztupinszki Z, Diossy M, Krzystanek M, Reiniger L, Csabai I, Favero F, et al. Migrating the SNP array-based homologous recombination deficiency measures to next generation sequencing data of breast cancer. NPJ breast cancer. 2018;4(1):1–4.

29. Nik-Zainal S, Davies H, Staaf J, Ramakrishna M, Glodzik D, Zou X, et al. Landscape of somatic mutations in 560 breast cancer whole-genome sequences. Nature. 2016;534(7605):47.

30. Newell F, Kong Y, Wilmott JS, Johansson PA, Ferguson PM, Cui C, et al. Whole-genome landscape of mucosal melanoma reveals diverse drivers and therapeutic targets. Nature communications. 2019;10(1):1–15.

31. Kane DP, Shcherbakova PV. A common cancer-associated DNA polymerase epsilon mutation causes an exceptionally strong mutator phenotype, indicating fidelity defects distinct from loss of proofreading. Cancer Res. 2014;74(7):1895–901.

32. Kahn RM, Gordhandas S, Maddy BP, Baltich Nelson B, Askin G, Christos PJ, et al. Universal endometrial cancer tumor typing: How much has immunohistochemistry, microsatellite instability, and MLH1 methylation improved the diagnosis of Lynch syndrome across the population? Cancer. 2019;125(18):3172–83.

33. Murai J, Huang SY, Renaud A, Zhang Y, Ji J, Takeda S, et al. Stereospecific PARP trapping by BMN 673 and comparison with olaparib and rucaparib. Mol Cancer Ther. 2014;13(2):433–43.

34. Mills GB, Timms KM, Reid JE, Gutin A, Krivak TC, Hennessy B, et al. Homologous recombination deficiency score shows superior association with outcome compared with its individual score components in platinum-treated serous ovarian cancer. Gynecologic Oncology. 2016;141:2–3.

35. Telli ML, Timms KM, Reid J, Hennessy B, Mills GB, Jensen KC, et al. Homologous recombination deficiency (HRD) score predicts response to platinum-containing neoadjuvant chemotherapy in patients with triple-negative breast cancer. Clinical cancer research. 2016;22(15):3764–73.

36. Polak P, Kim J, Braunstein LZ, Karlic R, Haradhavala NJ, Tiao G, et al. A mutational signature reveals alterations underlying deficient homologous recombination repair in breast cancer. Nature genetics. 2017.

37. Depreeuw J, Hermans E, Schrauwen S, Annibali D, Coenegrachts L, Thomas D, et al. Characterization of patient-derived tumor xenograft models of endometrial cancer for preclinical evaluation of targeted therapies. Gynecologic oncology. 2015;139(1):118–26.

38. Cuppens T, Depreeuw J, Annibali D, Thomas D, Hermans E, Gommé E, et al. Establishment and characterization of uterine sarcoma and carcinosarcoma patient-derived xenograft models. Gynecologic Oncology. 2017;146(3):538–45.

39. Ben-David U, Ha G, Tseng Y-Y, Greenwald NF, Oh C, Shih J, et al. Patient-derived xenografts undergo mouse-specific tumor evolution. Nature genetics. 2017;49(11):1567.

40. Butler KA, Hou X, Becker MA, Zanfagnin V, Enderica-Gonzalez S, Visscher D, et al. Prevention of human lymphoproliferative tumor formation in ovarian cancer patient-derived xenografts. Neoplasia. 2017;19(8):628–36.

41. Gibson WJ, Hoivik EA, Halle MK, Taylor-Weiner A, Cherniack AD, Berg A, et al. The genomic landscape and evolution of endometrial carcinoma progression and abdominopelvic metastasis. Nature genetics. 2016;48(8):848–55.

42. de la Vega LL, Samaha MC, Hu K, Bick NR, Siddiqui J, Hovelson DH, et al. Multiclonality and marked branched evolution of low-grade endometrioid endometrial carcinoma. Molecular Cancer Research. 2019;17(3):731–40.

43. Misale S, Yaeger R, Hobor S, Scala E, Janakiraman M, Liska D, et al. Emergence of KRAS mutations and acquired resistance to anti-EGFR therapy in colorectal cancer. Nature. 2012;486(7404):532–6.

44. Pao W, Wang TY, Riely GJ, Miller VA, Pan Q, Ladanyi M, et al. KRAS mutations and primary resistance of lung adenocarcinomas to gefitinib or erlotinib. PLoS medicine. 2005;2(1):e17.

45. Priestley P, Baber J, Lolkema MP, Steeghs N, de Bruijn E, Shale C, et al. Pan-cancer whole-genome analyses of metastatic solid tumours. Nature. 2019:1–7.

46. Bielski CM, Zehir A, Penson AV, Donoghue MT, Chatila W, Armenia J, et al. Genome doubling shapes the evolution and prognosis of advanced cancers. Nature genetics. 2018;50(8):1189–95.

47. Nones K, Johnson J, Newell F, Patch A, Thorne H, Kazakoff S, et al. Whole-genome sequencing reveals clinically relevant insights into the aetiology of familial breast cancers. Annals of Oncology. 2019;30(7):1071–9.

48. Maura F, Degasperi A, Nadeu F, Leongamornlert D, Davies H, Moore L, et al. A practical guide for mutational signature analysis in hematological malignancies. Nature communications. 2019;10(1):1–12.

49. Sanij E, Hannan KM, Xuan J, Yan S, Ahern JE, Trigos AS, et al. CX-5461 activates the DNA damage response and demonstrates therapeutic efficacy in high-grade serous ovarian cancer. Nature communications. 2020;11(1):1–18.

50. Shen J, Zhao W, Ju Z, Wang L, Peng Y, Labrie M, et al. PARPi triggers the STING-dependent immune response and enhances the therapeutic efficacy of immune checkpoint blockade independent of BRCAness. Cancer research. 2019;79(2):311–9.

