## Additional File 1 for "Genomic analysis of patient-derived xenograft models reveals intra-tumor heterogeneity in endometrial cancer and can predict tumor growth inhibition with talazoparib"

**Bonazzi et al.**

### Supplementary Materials

#### Supplementary Methods

*SNP Array Analysis*

Single nucleotide polymorphism (SNP) arrays were scanned on an iScan (Illumina). Data was processed using the Genotyping module (v.1.9.4) in GenomeStudio v.2011.1 (Illumina) to calculate B-allele frequencies (BAF) and logR ratios. GAP (1) was used to call somatic regions of copy number change after low-quality probes assessed in the matched normal sample with GenCall (GC) score of <0.7 were removed.

*Alignment to human and mouse genome references*

Reads were aligned to a combined human/mouse (GRCh37/GRCm38 Nodshiltj background) reference using BWA-mem (v0.7.15). Only read paired aligned to the human sequences with a mapping quality score (MAPQ) = 60 were used for downstream analysis (Table S8). We confirmed that all reads mapping to human reference with MAPQ60 had lower secondary alignment scores (XS) to mouse reference compared with primary alignment scores (AS).

*Variant calling*

A dual variant-calling strategy was used. Substitutions were detected using an in-house developed tool qSNP (2) and GATK Haplotype caller (3), and insertions or deletions (indels) of 1-60 bp in length were called with GATK Haplotype caller. Filters were applied to ensure only high confidence variants were reported (4). For somatic coding variants in WES samples, the variant filtering was less stringent to include variants with lower coverage. The less-stringent filters included: strand bias of alternate allele, less than 5 variant reads, variant also found in pileup of normal and homopolymer filter. Variants were annotated with Ensembl v75 gene feature information and transcript or protein consequences using SnpEff (v4.2) (5).

Large structural rearrangements were identified using an in-house developed tool qSV for WGS (4). Somatic copy number alterations (CNA), tumor ploidy and purity were determined with ascatNgs for WGS (6) and GAP for SNP arrays (1). Homozygous deletions (CN<1), deletions (CN<2) gains (CN≥ploidy + 1) and amplifications (CN≥ploidy +3) were considered in the analysis. Copy number alterations (CNA) and structural variant (SV) events were annotated against Ensembl. Percentage of genome with CNA was determined from SNP array and WGS CNA data for PDX tumour samples, as percentage of genome (autosomal chromosomes) with copy number ≠ 2.

*Heterogeneity analysis*

Somatic mutation comparison between primary and PDX tumour samples was performed for four mismatch-repair deficient (MMRd) models with WES data and three carcinosarcoma models with WGS. A union of all pass-filter missense variants was generated for each model, and then the genomic positions of these variants were interrogated with qBasePileup to capture any variants that may have been missed during variant calling, because of low frequency or other quality filters. The overlaps of pileup variants were visualised using eulerR package. The generated pileup list of variants with allele-specific copy number information, extracted from ascatNGS (6) or GAP (1) output (for WGS and SNP arrays, respectively), was then used to identify mutational clusters using PyClone (v0.13.1) (7) with pyclone_beta_binomial emission density, random seed set to 20 and thin parameter set to 10. The top five mutational clusters with the most variants were selected for determining clonal evolution using ClonEvol package (8).

Copy number clonality analysis was performed on the WGS data of tumor-normal pairs using Battenberg (v2.2.5) with default settings.

*MSI status*

The level of microsatellite instability (MSI) was assessed using MSIsensor (v0.2) on tumor-normal pairs of primary and PDX tumor samples using suggested parameters for WES and WGS data (9). Samples with an MSI score of >3 were classified as MSI-high.

*HRD score assessment*

Homologous recombination deficiency (HRD) scores were assessed on SNP array and WGS data using scarHRD (10) package on the allele-specific copy number information determined by either GAP (1) or ascatNGS (6).

*Signature analysis*

Mutational signature analysis was performed using two approaches with SigProfiler and deconstructSigs. *De novo* signatures were identified on somatic single nucleotide variants from WES data using non-negative matrix factorization (NMF) with SigProfiler, as previously described (11). The identified signatures were then compared to the 30 known COSMIC v2 signatures. The optimal number of signatures was chosen based on a number of parameters: stability, reconstruction error and cosine similarities to COSMIC signatures. The contribution of each *de novo* signature to a sample's mutational profile was assigned using SignatureEstimation package (12). To determine the potential relative contribution of signature 3 (HRD-associated signature), deconstructSigs package (13) was used to estimate the contribution of the sample's mutations to the full catalogue of COSMIC v2 mutational signatures, using default setting. Minimum mutation signature contribution cut-off was set to 15%.

SV signatures were identified using the same approach as used for *de novo* mutational signature analysis. SV events were classified into 32 previously defined categories based on event type, size and breakpoint clustering (14). SV signature analysis was performed as previously described (15).

*Targetable Mutations Analysis*

Cancer Genome Interpreter analysis was performed in July 2019 to identify potentially targetable mutations in the PDX models. Short somatic variants, homozygous deletions and amplifications in coding regions observed in all PDX samples were used as input. Only known pathogenic variants and predicted drivers (tier 1, 2) were considered as biomarkers. Biomarkers were matched with drugs using the Cancer Biomarker Database within Cancer Genome Interpreter. The drug prescription output from Cancer Genome Interpreter was filtered to select variants that were "complete" alterations (alterations that match the specific amino acid change in the gene which constitutes an actionable variant for a specific drug). Only drug sensitivity biomarkers with FDA approved drugs or drugs currently in clinical trials were included. Chemotherapy and steroid compounds were not included.

#### Supplementary Tables

**Table S1. Clinical and histopathological characteristics of patient-derived xenograft (PDX) models of uterine cancers.**

| PDX # | Age | Stage | Grade | Histology | Chemotherapy response | Patient cancer status | Current Mortality status |
|---|---|---|---|---|---|---|---|
| PDX03 | 60 | Ic | 3 | UCS | Chemoradiation | Recurred | DWD |
| PDX12 | 70 | Ia | 3 | HG EEC | Radiation only | - | NED |
| PDX21 | 67 | IIIc | 3 | HG EEC | Radiation only | Recurred | DWD |
| PDX23 | 58 | IIIb | 3 | HG SEC | Chemoradiation | Recurred | DWD |
| PDX24 | 86 | Ib | 3 | HG EEC | Radiation only | - | NED |
| PDX49 | 70 | Ia | 3 | UCS | Chemoradiation | - | NED |
| PDX52 | 83 | Ib | 2 | LG EEC | Radiation only | Recurred | DWD |
| PDX53 | 43 | II | 3 | HG SEC | Chemoradiation | - | NED |
| PDX56 | 79 | Ia | 3 | UCS | Chemoradiation | Recurred | AWD |
| PDX58 | 62 | Ib | 3 | HG EEC | No treatment | Recurred | DWD |
| PDX59 | 71 | Ib | 3 | HG EEC | Radiation only | - | NED |

UCS — Uterine Carcinosarcoma; HG EEC — high-grade endometrioid endometrial cancer, HG SEC — high-grade serous endometrial cancer; LG EEC — low-grade endometrioid endometrial cancer; DWD — died with disease; AWD — Alive with disease; NED — alive with no evidence of disease.

**Table S2. Successful PDX engraftment by grade and storage conditions.**

| Tumor tissue condition | G1 | G2 | G3 | Total EC PDXs |
|---|---|---|---|---|
| Fresh | 0/8 | 2/6 (33%) | 13/19 (68%) | 15/33 (45%) |
| Overnight at 4oC | 1/4 (25%) | | 1/5 (20%) | 2/9 (22%) |
| Viably Frozen (-80oC) | 0/2 | 0/5 | 1/4 (25%) | 1/11 (9%) |

**Table S3. Genes with common SNV and short indels or CNA events identified by TCGA UCEC and TCGA UCS studies.**

| Gene | Type of somatic event | TCGA study |
|---|---|---|
| *ARHGAP35* | SNV and short indels | UCS |
| *ARID1A* | SNV and short indels | UCEC |
| *ARID5B* | SNV and short indels | UCEC |
| *CDH4* | SNV and short indels | UCS |
| *CTNNB1* | SNV and short indels | UCEC |
| *FBXW7* | SNV and short indels | UCEC, UCS |
| *KRAS* | SNV and short indels | UCEC, UCS |
| *PIK3CA* | SNV and short indels | UCEC, UCS |
| *PIK3R1* | SNV and short indels | UCEC, UCS |
| *POLE* | SNV and short indels | UCEC |
| *PPP2R1A* | SNV and short indels | UCEC, UCS |
| *PTEN* | SNV and short indels | UCEC, UCS |
| *RB1* | SNV and short indels | UCS |
| *RPL22* | SNV and short indels | UCEC |
| *SPOP* | SNV and short indels | UCS |
| *TP53* | SNV and short indels | UCEC, UCS |
| *U2AF* | SNV and short indels | UCS |
| *ZBTB7B* | SNV and short indels | UCS |
| *BCL2L1* | CNA | UCS |
| *CCNE1* | CNA | UCEC, UCS |
| *ERBB2* | CNA | UCEC, UCS |
| *FGFR1* | CNA | UCEC |
| *FGFR3* | CNA | UCEC, UCS |
| *IGF1R* | CNA | UCEC |
| *KAT6A* | CNA | UCS |
| *LRP1B* | CNA | UCEC |
| *MDM2* | CNA | UCS |
| *MYC* | CNA | UCEC, UCS |
| *RIT1* | CNA | UCS |
| *SOX17* | CNA | UCEC |
| *TERC* | CNA | UCS |

TCGA — The Cancer Genome Atlas; UCEC — Uterine corpus endometrial carcinoma; UCS — uterine carcinosarcoma.

**Table S4. Somatic coding variants and CNAs detected in PDX tumour samples in genes relevant to endometrial cancer.**

These variants are summarized in Fig 2. PDX03 and PDX49 models were analyzed with WES and WGS, where WES samples are denoted with *. Attached as separate document.

**Table S5. HR DNA repair associated genes.**

| Gene | RefSeq transcript | HR-related gene |
|---|---|---|
| *ATM* | NM_000051.3 | yes |
| *ATR* | NM_001184.3 | |
| *BARD1* | NM_000465.3 | yes |
| *BRCA1* | NM_007294.3 | yes |
| *BRCA2* | NM_000059.3 | yes |
| *BRIP1* | NM_032043.2 | yes |
| *CDK12* | NM_016507.3 | yes |
| *CHEK1* | NM_001114121.2 | yes |
| *CHEK2* | NM_007194.3 | yes |
| *FANCA* | NM_000135.2 | |
| *FANCB* | NM_001018113.2 | |
| *FANCC* | NM_000136.2 | |
| *FANCD2* | NM_033084.4 | |
| *FANCE* | NM_021922.2 | |
| *FANCF* | NM_022725.3 | |
| *FANCG* | NM_004629.1 | |
| *FANCI* | NM_001113378.1 | |
| *FANCL* | NM_001114636.1 | |
| *FANCM* | NM_020937.3 | |
| *MRE11* | NM_005591.3 | yes |
| *NBN* | NM_002485.4 | yes |
| *PALB2* | NM_024675.3 | yes |
| *RAD50* | NM_005732.4 | yes |
| *RAD51* | NM_002875.4 | yes |
| *RAD51B* | NM_133509.3 | yes |
| *RAD51C* | NM_058216.2 | yes |
| *RAD51D* | NM_002878.3 | yes |

**Table S6. HR DNA repair gene variants detected in EC PDX models.**
Attached as separate document.

**Table S7. HR DNA repair gene variants detected in TCGA-UCEC and TCGA-UCS studies.**
Attached as separate document.

**Table S8. Sequencing read alignment statistics for human and mouse genome references for WES and WGS samples.**
Attached as separate document.

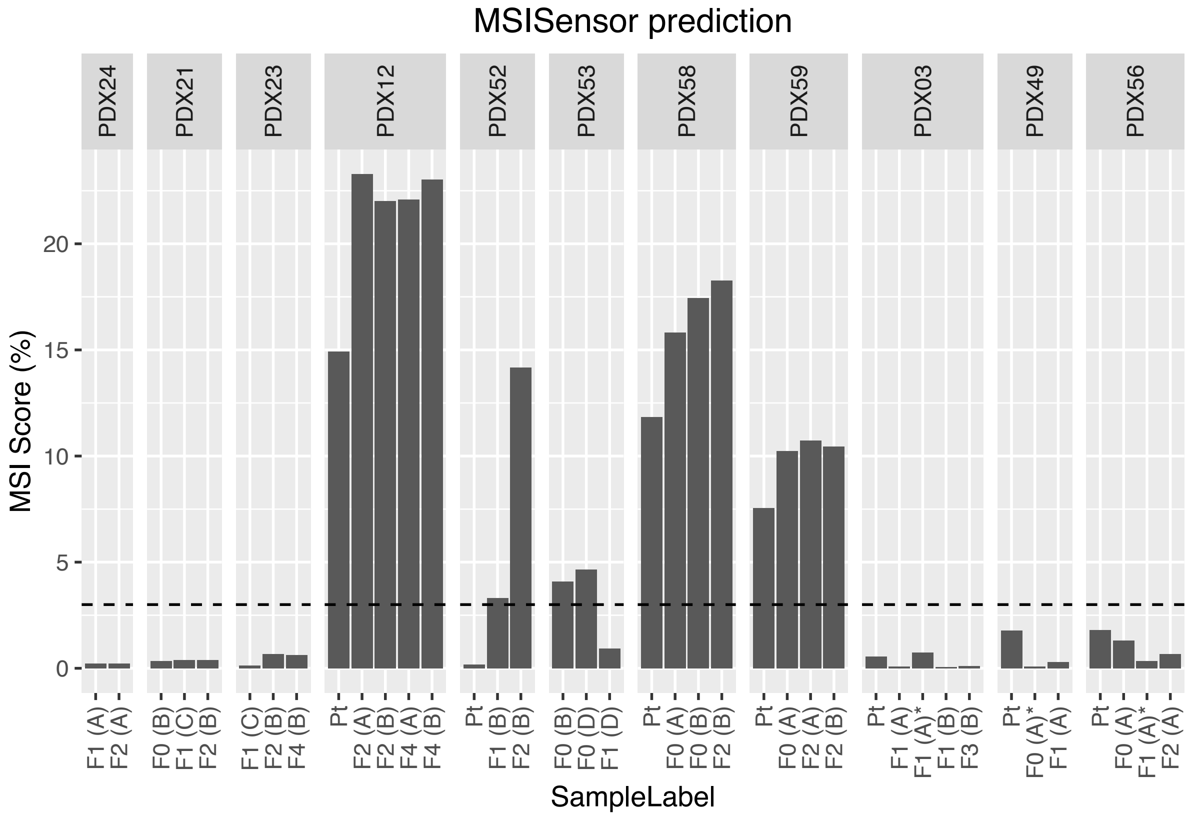


**Fig. S1. MSI score determined by MSIsensor in primary and PDX tumour samples assessed by WES.** Tumour samples are grouped by patient ID. PDX samples are labelled by passage number and lineage in brackets. PDX03 and PDX49 were analyzed with WES and WGS, with WES samples denoted with *. The MSI cut-off of 3 is shown with the horizontal dash line. MSI — microsatellite instability; PDX — patient-derived xenograft; WES — whole-exome sequencing; Pt — patient.

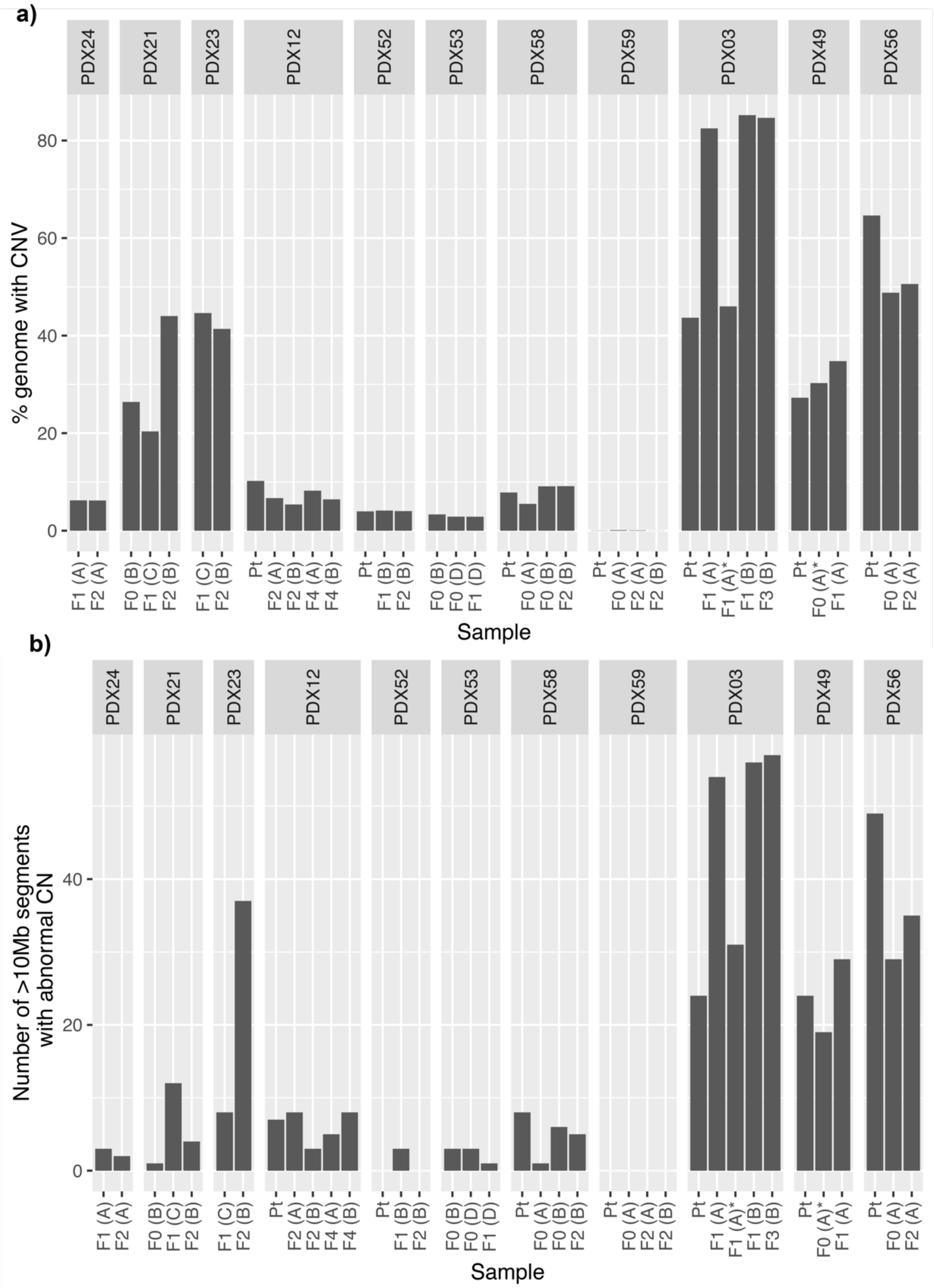


**Fig. S2. Degree of copy number instability observed in primary and PDX tumour samples assessed by: a** Percentage of genome with CNA for autosomal chromosomes only; and **b** Number of abnormal CN segments, including CN neutral LOH, of >10Mb. The CNA were determined from SNP arrays, except for PDX56, where WGS CNA data was used. Tumour samples are grouped by patient ID PDX03 and PDX49 were analyzed with WES and WGS, with WES samples denoted with *. PDX samples are labelled by passage number and lineage in brackets. CN — copy number; CNA — CN alterations; LOH — loss of heterozygosity; WGS — whole-genome sequencing.

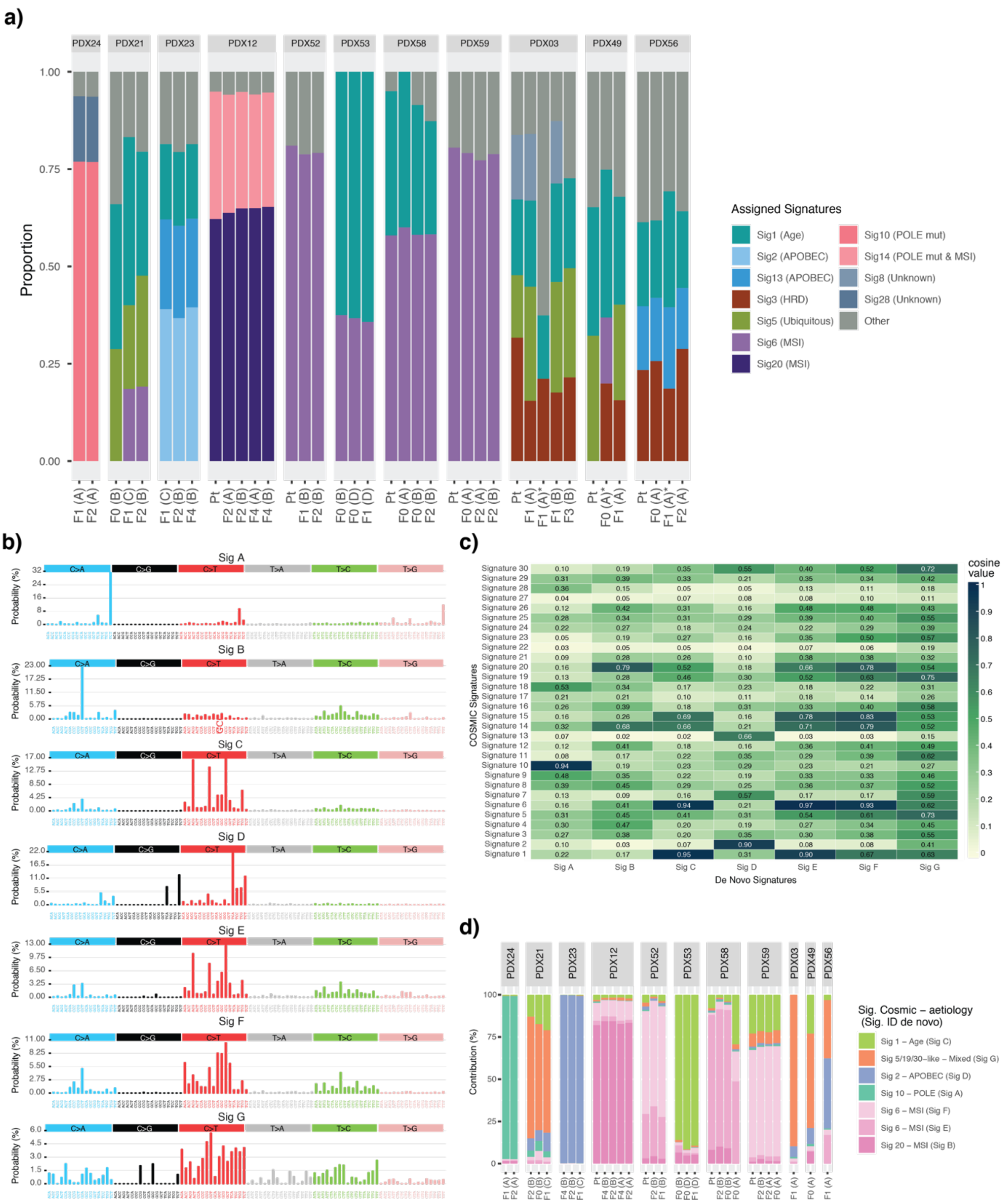


**Fig. S3. Mutational signature analysis in PDX models. a** Mutational signature analysis using deconstructSigs. Minimum signature contribution cut-off of 15% was used to avoid signature overfitting. PDX03 and PDX49 were analyzed with WES and WGS, with WES samples denoted with *. **B** The mutational type probability for each substitution in a trinucleotide context of seven *de novo* signatures identified by SigProfiler. **c** Cosine similarity matrix of seven de novo signatures and 30 known COSMIC (v2) signatures. **d** The relative contribution of de novo signatures to the mutational profile of each tumour sample analyzed using WES. Tumour samples are grouped by patient ID. PDX samples are labelled by passage number and lineage in brackets. Pt — patient.

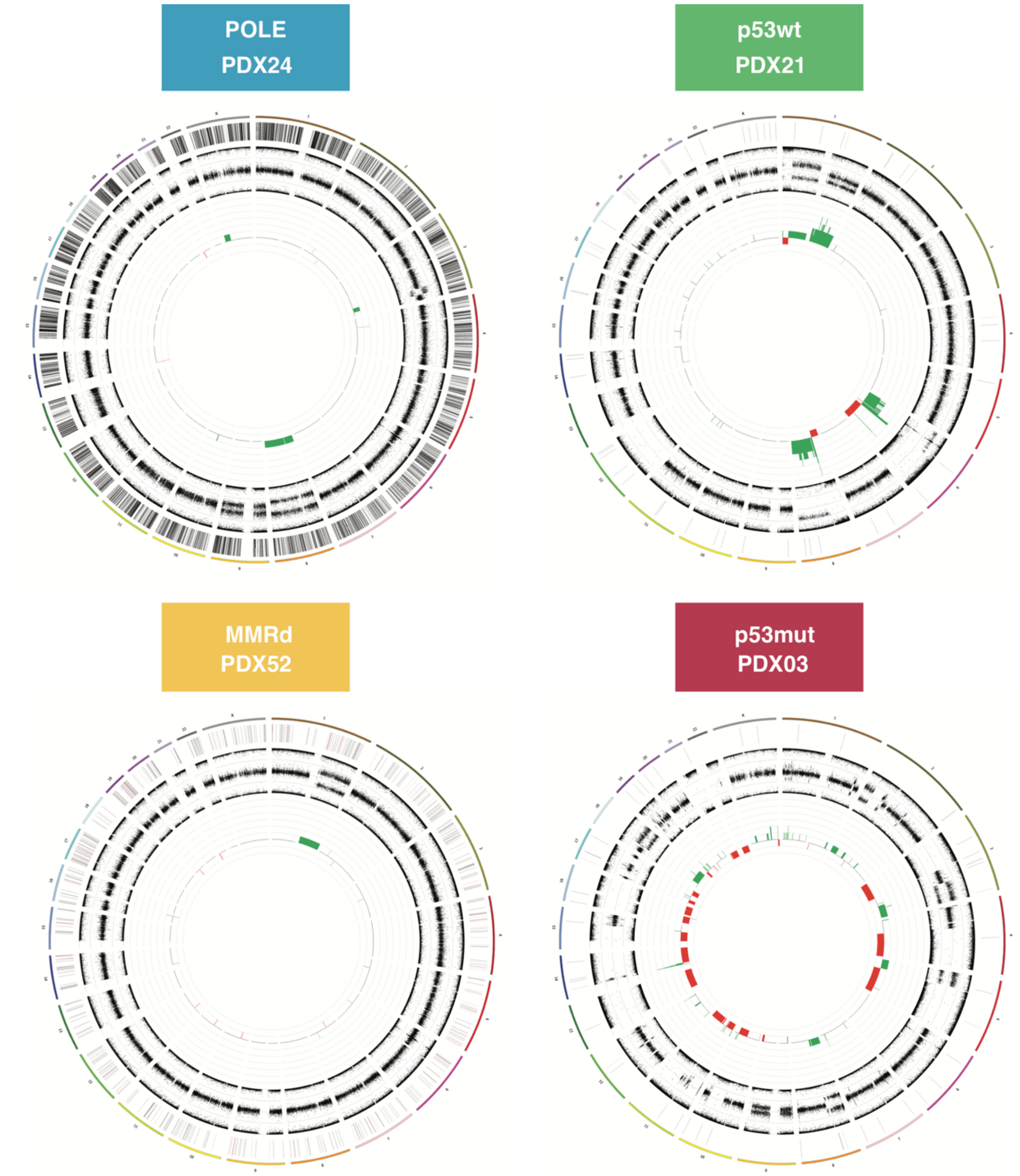


**Fig. S4. Circos plots of representative PDX samples show genome-wide characteristics of each subtype.** The outer ring shows chromosomes 1-22 and X. The next ring shows somatic coding variants, where substitutions are colored in black and indels are in red. The next ring shows BAFs determined by SNP arrays. The most inner ring shows copy number segments identified by GAP from SNP array data, where gains are colored in green and deletions are in red. BAF — B-allele frequency.

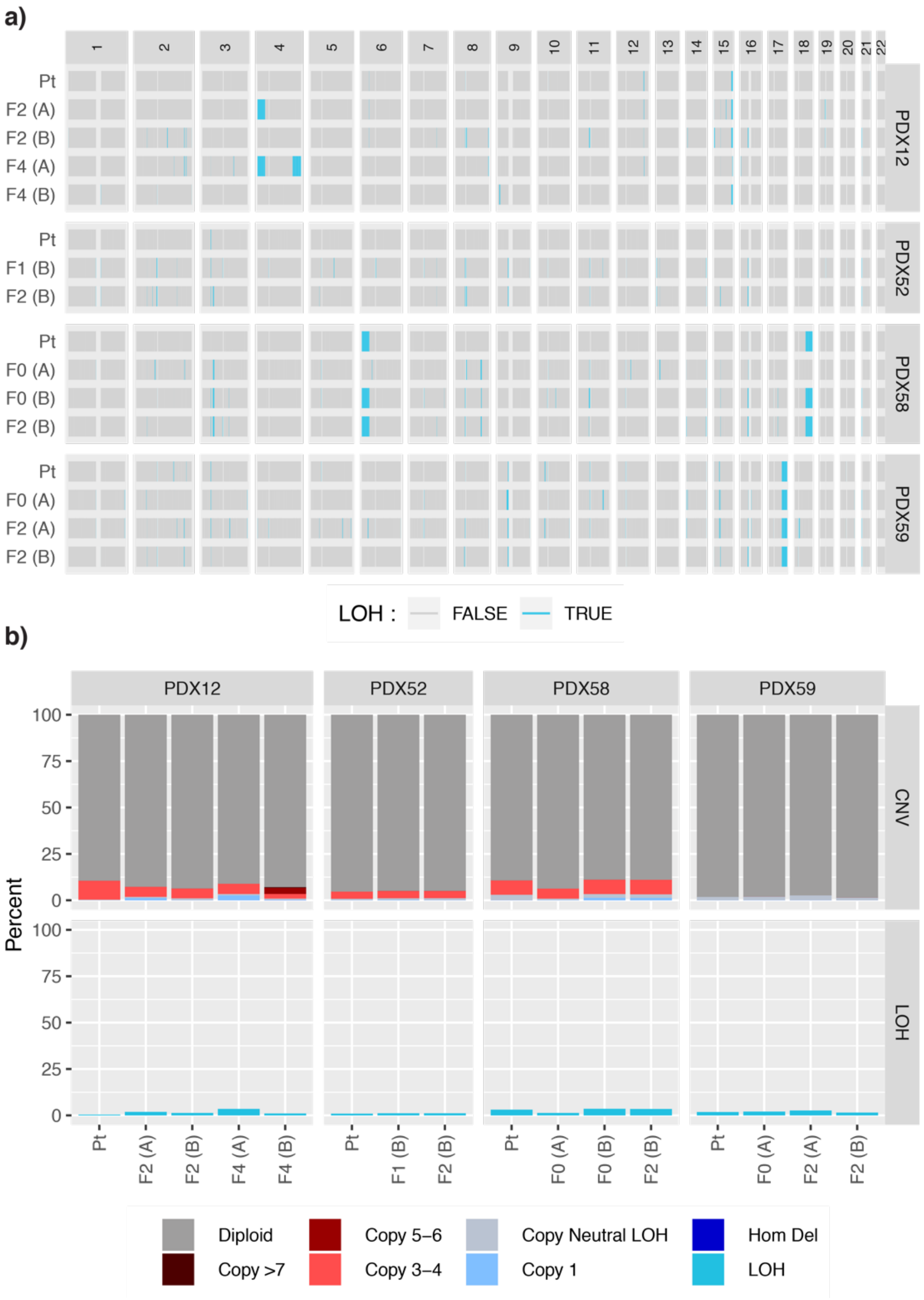


**Fig. S5. Genome-wide CNA profiles observed in the primary and matched PDX samples of the MMRd EC PDX models: a** LOH profiles analyzed by SNP arrays; **b** genome-wide summary of CNA and LOH. Tumour samples are grouped by patient ID. PDX samples are labelled by passage number and lineage in brackets.

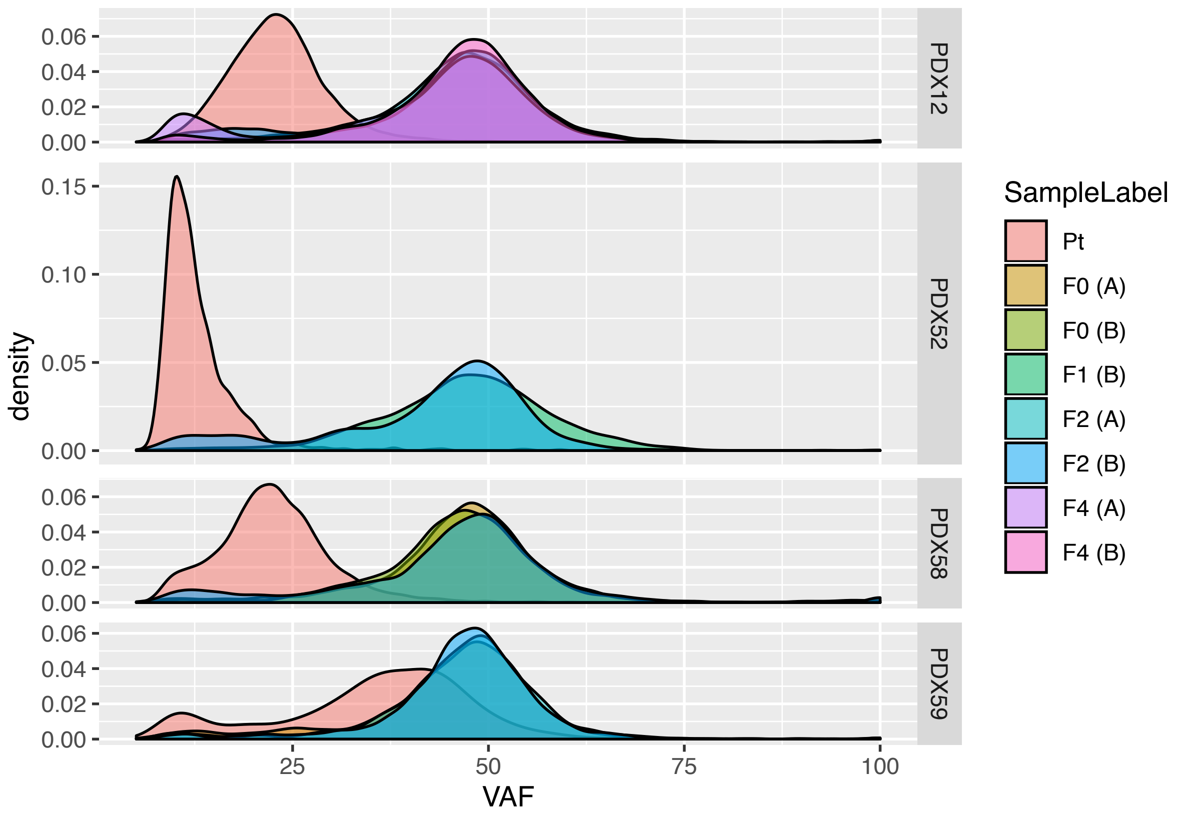


**Fig. S6. Lower tumour purity observed in primary tumour samples compared to matched PDX samples as assessed by somatic variant allele frequency.** Variant allele frequency of all somatic variants detected by WES is shown. Tumour samples are grouped by patient ID. PDX samples are labelled by passage number and lineage in brackets.

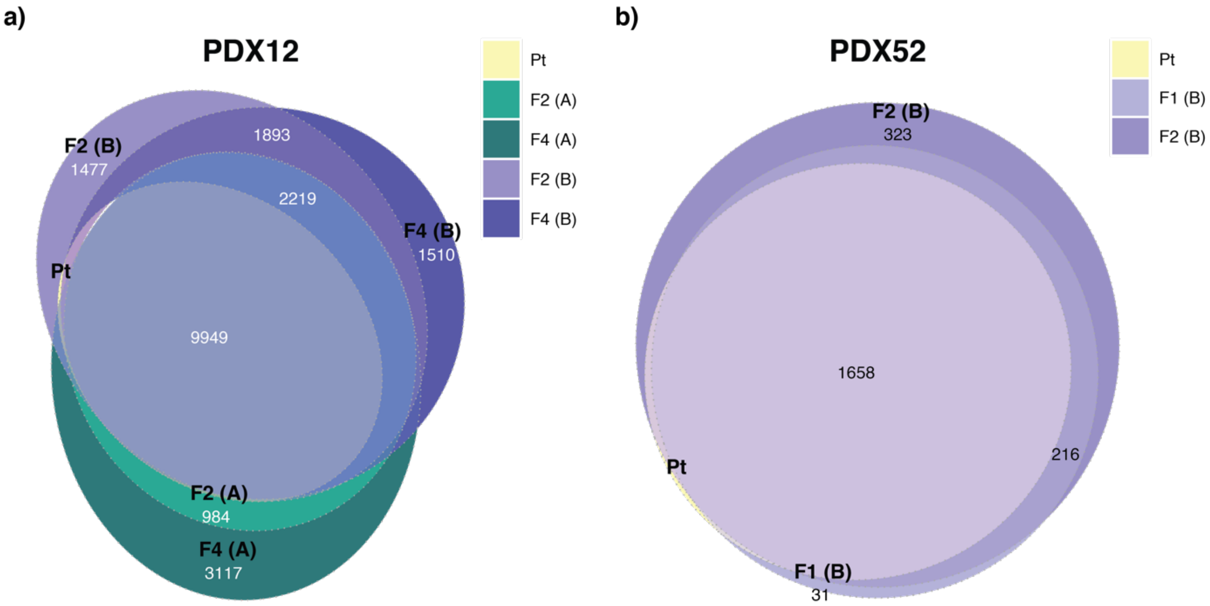


**Fig. S7. Mutational heterogeneity visualized by Euler diagrams.** Somatic substitutions called by qBasepileup in **a** PDX12 and **b** PDX52 mismatch repair deficient models. PDX — patient-derived xenograft.

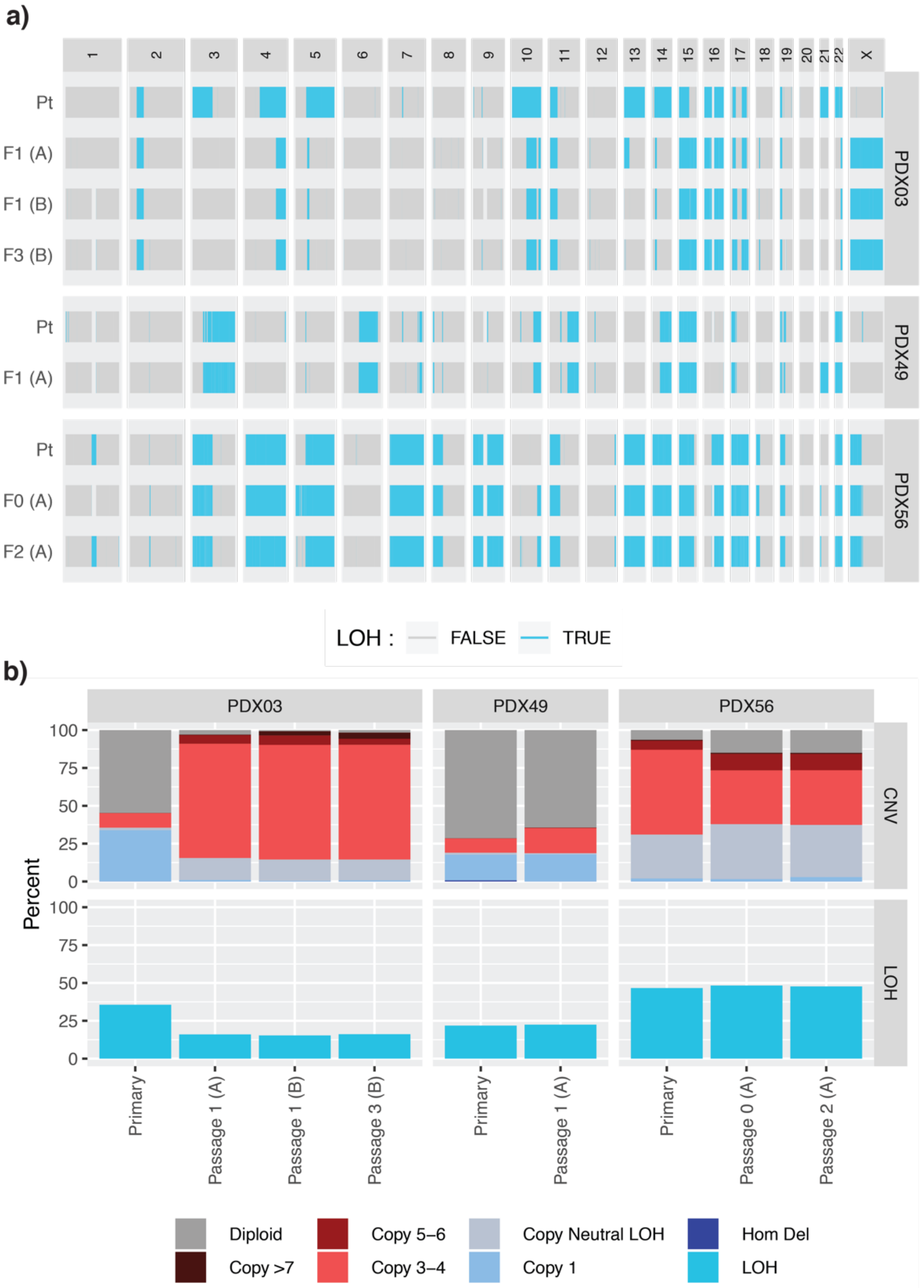


**Fig. S8. Genome-wide CNA profiles observed in the primary and matched PDX samples of the p53mut UCS PDX models: a** LOH profiles analyzed by SNP arrays; **b** Genome-wide summary of CNA and LOH.

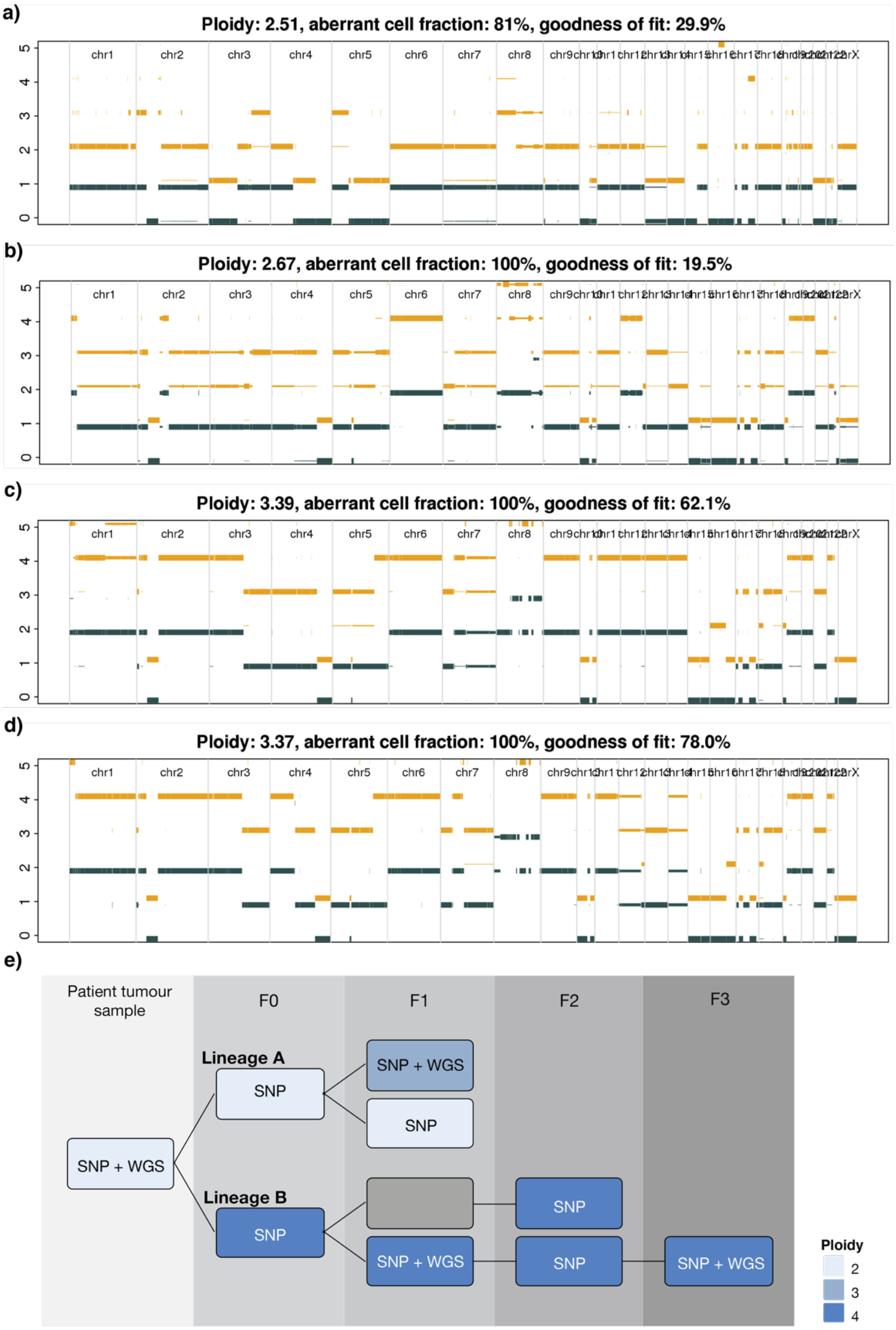


**Fig. S9. CNA profiles of the primary and matched PDX samples of p53mut carcinosarcoma PDX03 model.** Battenberg subclonal CN profiles for **a** primary tumour, **b** Passage 1 — Lineage A sample, **c** Passage 1 — Lineage B sample and **d** Passage 3 — Lineage B sample. Thick lines show the predominant CN state, thin lines show subclonal CN, yellow color denotes major allele CN, and dark grey denotes minor allele CN. PDX — patient-derived xenograft; CN — copy number. **e** Ploidy estimations determined from SNP array or WGS data, where it was available.

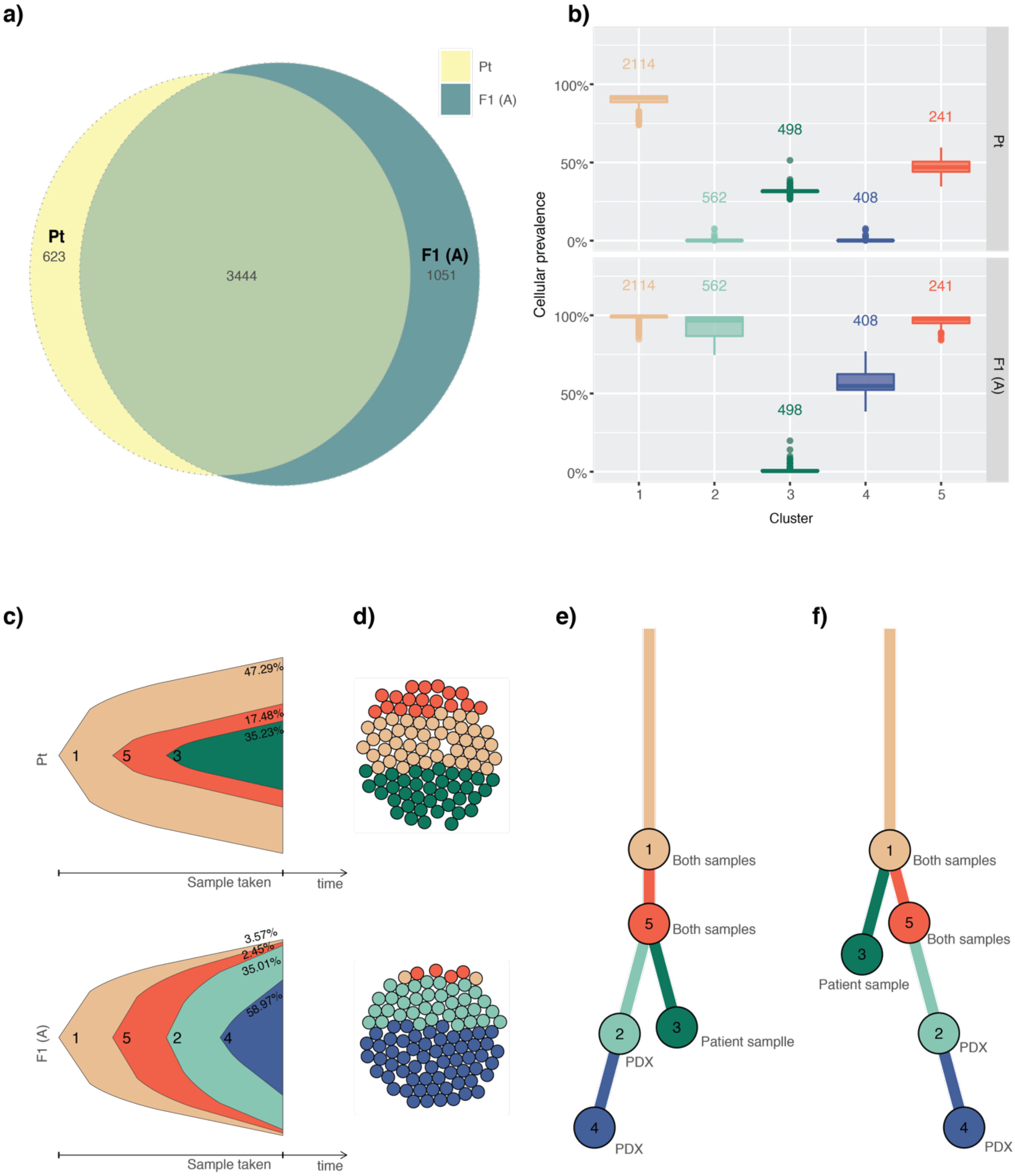


**Fig. S10. Clonal evolution analysis of mutations in the p53mut carcinosarcoma PDX49 model. a** Mutational heterogeneity visualized by euler diagrams of somatic substitutions called by qBasepileup. **b** Cellular prevalence of the top five mutational clusters with ≥5% of all somatic substitutions detected by PyClone. Values shown above boxplots represent the number of mutations contributing to each cluster. **c** Fish plots and **d** cellular population depictions of the top five mutational clusters. Percentages shown in the fish plots are the estimated proportions of cells containing that mutational cluster. **e** The selected clonal evolution tree inferred by ClonEvol. **f** An alternative solution for the clonal evolution tree by ClonEvol.

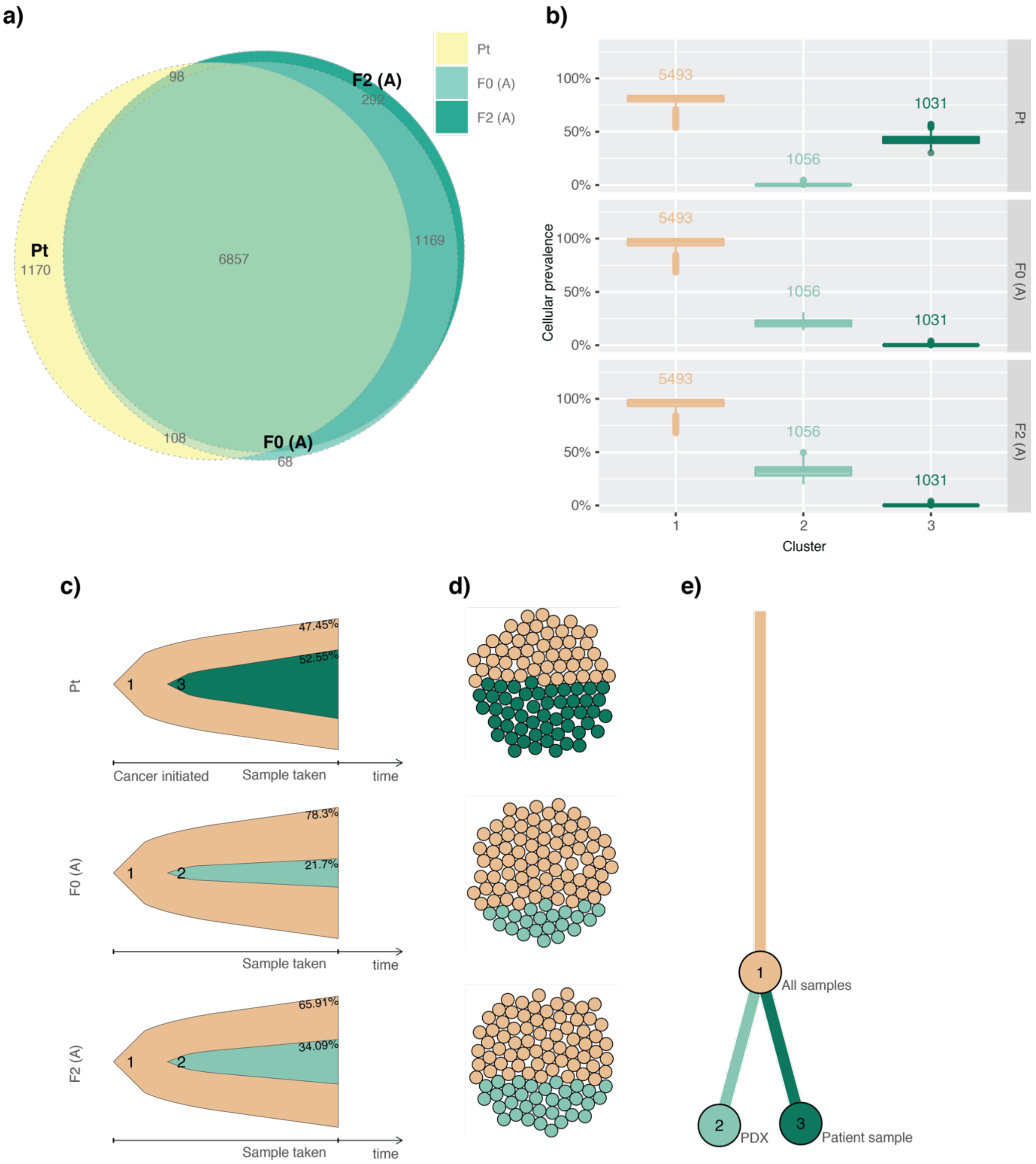


**Fig. S11. Clonal evolution analysis of mutations in the p53mut carcinosarcoma PDX56 model. a** Mutational heterogeneity visualized by euler diagrams of somatic substitutions called by qBasepileup. **b** Cellular prevalence of the top three mutational clusters with ≥5% of all somatic substitutions detected by PyClone. Values shown above boxplots represent the number of mutations contributing to each cluster. **c** Fish plots and **d** cellular population depictions of the top three mutational clusters. Percentages shown in the fish plots are the estimated proportions of cells containing that mutational cluster. **e** The clonal evolution tree inferred by ClonEvol.

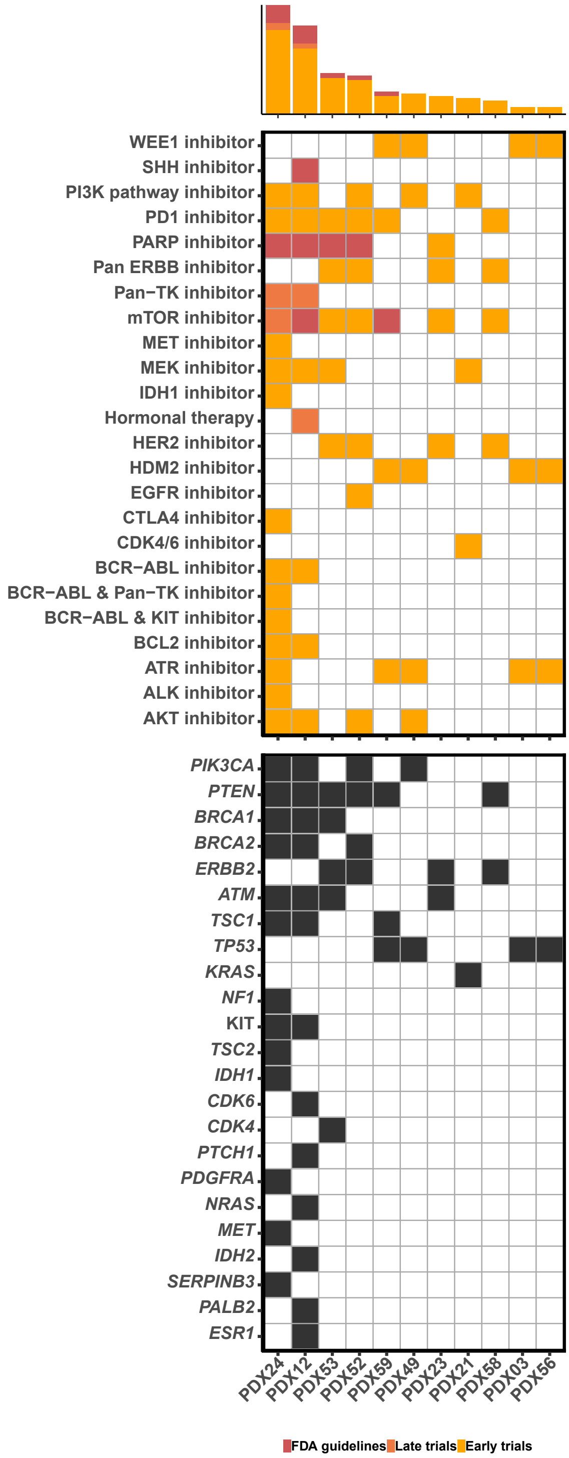


**Fig. S12. Potentially actionable genomic alterations in PDX models identified using Cancer Genome Interpreter.** Genomic alterations are shown in black in the bottom panel.

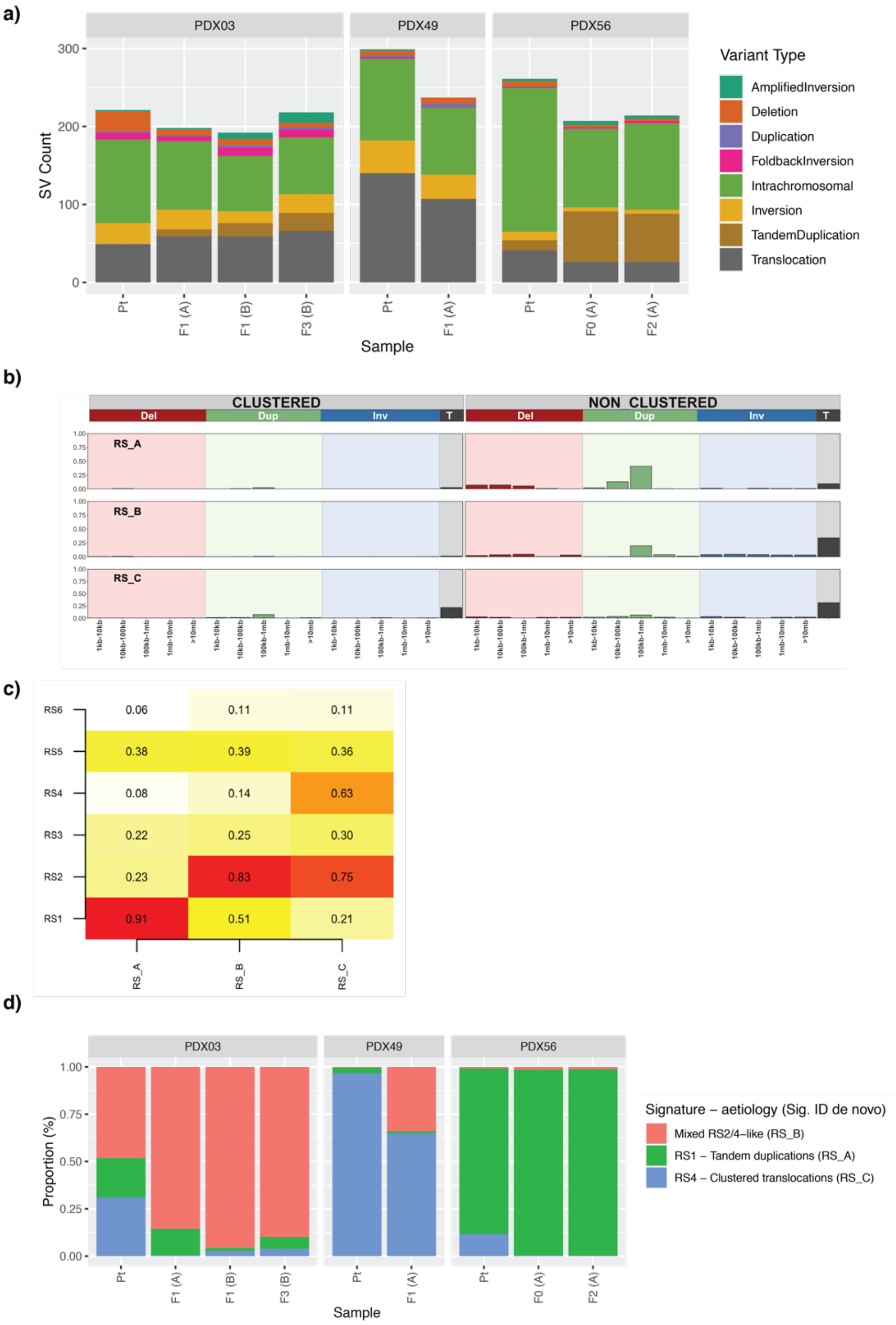


**Fig. S13. De novo rearrangement signature analysis in the three p53mut carcinosarcoma models assessed by WGS. a** Structural variant types detected in the PDX models that were used for rearrangement signature analysis. **b** The rearrangement class profile of the three de novo rearrangement signatures. Rearrangements were classified into 32 categories based on the rearrangement size, type and whether breakpoints are clustered or non-clustered. **c** Cosine similarity matrix of the three de novo rearrangement signatures and previously described rearrangement signatures in breast cancer. **d** The relative contribution of de novo signatures to the mutational profile of each tumour sample. Tumour samples are grouped by patient ID. PDX samples are labelled by passage number and lineage in brackets. PDX — patient-derived xenograft; WGS — whole-genome sequencing; Del — deletion, Dup — duplication, Inv — inversion, T — translocation.

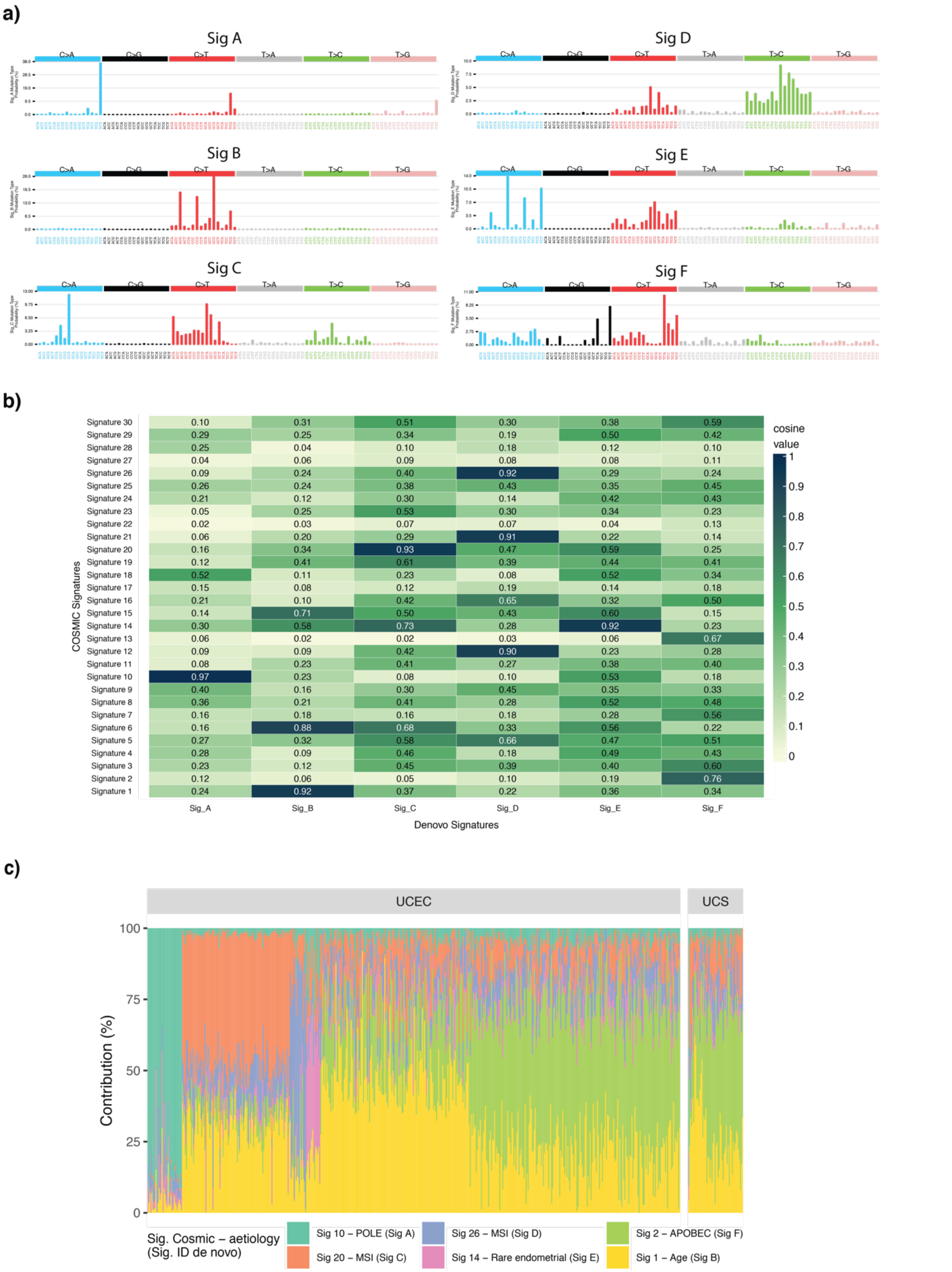


**Fig. S14. De novo mutational signature analysis in TCGA UCEC and UCS cohorts. a** The mutational type probability for each substitution in a trinucleotide context of six de novo signatures identified by SigProfiler. **b** Cosine similarity matrix of six de novo signatures and 30 known COSMIC (v2) signatures. **c** The relative contribution of de novo signatures to the mutational profile of each tumour sample.

#### References


1. Popova T, Manié E, Stoppa-Lyonnet D, Rigaill G, Barillot E, Stern MH. Genome Alteration Print (GAP): a tool to visualize and mine complex cancer genomic profiles obtained by SNP arrays. Genome biology. 2009;10(11):1-14.
2. Kassahn KS, Holmes O, Nones K, Patch AM, Miller DK, Christ AN, et al. Somatic point mutation calling in low cellularity tumors. PLoS One. 2013;8(11):e74380.
3. McKenna A, Hanna M, Banks E, Sivachenko A, Cibulskis K, Kernytsky A, et al. The Genome Analysis Toolkit: a MapReduce framework for analyzing next-generation DNA sequencing data. Genome Res. 2010;20(9):1297-303.
4. Patch AM, Christie EL, Etemadmoghadam D, Garsed DW, George J, Fereday S, et al. Whole-genome characterization of chemoresistant ovarian cancer. Nature. 2015;521(7553):489-94.
5. Cingolani P, Platts A, Wang LL, Coon M, Nguyen T, Wang L, et al. A program for annotating and predicting the effects of single nucleotide polymorphisms, SnpEff: SNPs in the genome of Drosophila melanogaster strain w1118; iso-2; iso-3. Fly. 2012;6(2):80-92.
6. Raine KM, Van Loo P, Wedge DC, Jones D, Menzies A, Butler AP, et al. ascatNgs: Identifying Somatically Acquired Copy-Number Alterations from Whole-Genome Sequencing Data. Current protocols in bioinformatics. 2016;56(1):15.9. 1-.9. 7.
7. Roth A, Khattra J, Yap D, Wan A, Laks E, Biele J, et al. PyClone: statistical inference of clonal population structure in cancer. Nature methods. 2014;11(4):396-8.
8. Dang H, White B, Foltz S, Miller C, Luo J, Fields R, et al. ClonEvol: clonal ordering and visualization in cancer sequencing. Annals of oncology. 2017;28(12):3076-82.
9. Niu B, Ye K, Zhang Q, Lu C, Xie M, McLellan MD, et al. MSIsensor: microsatellite instability detection using paired tumor-normal sequence data. Bioinformatics. 2014;30(7):1015-6.
10. Sztupinszki Z, Diossy M, Krzystanek M, Reiniger L, Csabai I, Favero F, et al. Migrating the SNP array-based homologous recombination deficiency measures to next generation sequencing data of breast cancer. NPJ breast cancer. 2018;4(1):1-4.
11. Alexandrov LB, Nik-Zainal S, Wedge DC, Aparicio SA, Behjati S, Biankin AV, et al. Signatures of mutational processes in human cancer. Nature. 2013;500(7463):415-21.
12. Huang X, Wojtowicz D, Przytycka TM. Detecting presence of mutational signatures in cancer with confidence. Bioinformatics. 2018;34(2):330-7.
13. Rosenthal R, McGranahan N, Herrero J, Taylor BS, Swanton C. DeconstructSigs: delineating mutational processes in single tumors distinguishes DNA repair deficiencies and patterns of carcinoma evolution. Genome biology. 2016;17(1):1-11.
14. Nik-Zainal S, Davies H, Staaf J, Ramakrishna M, Glodzik D, Zou X, et al. Landscape of somatic mutations in 560 breast cancer whole-genome sequences. Nature. 2016;534(7605):47.
15. Newell F, Kong Y, Wilmott JS, Johansson PA, Ferguson PM, Cui C, et al. Whole-genome landscape of mucosal melanoma reveals diverse drivers and therapeutic targets. Nature communications. 2019;10(1):1-15.
